# Sigma receptor ligands are potent anti-prion compounds that act independently of sigma receptor binding

**DOI:** 10.1101/2023.11.28.569035

**Authors:** Robert C. C. Mercer, Nhat T. T. Le, Mei C. Q. Houser, Aaron B. Beeler, David A. Harris

## Abstract

Prion diseases are invariably fatal neurodegenerative diseases of humans and other animals for which there are no treatment options. Previous work from our laboratory identified phenethyl piperidines as novel class of anti-prion compounds. While working to identify the molecular target(s) of these molecules, we unexpectedly discovered ten novel anti-prion compounds based on their known ability to bind to the sigma receptors, σ_1_R and _2_R, which are currently being tested as therapeutic or diagnostic targets for cancer and neuropsychiatric disorders. Surprisingly, however, knockout of the respective genes encoding σ_1_R and σ_2_R (*Sigmar1* and *Tmem97*), in prion infected N2a cells did not alter the anti-prion activity of these compounds, demonstrating that these receptors are not the direct targets responsible the anti-prion effects of their ligands. Further investigation of the most potent molecules established that they are efficacious against multiple prion strains and protect against downstream prion-mediated synaptotoxicity. While the precise details of the mechanism of action of these molecules remains to be determined, the present work forms the basis for further investigations of these compounds in pre-clinical studies. Given the therapeutic utility of several of the tested compounds, including rimcazole and haloperidol for neuropsychiatric conditions, (+)-pentazocine for neuropathic pain, and the ongoing clinical trials of SA 4503 and ANAVEX2-73 for ischemic stroke and Alzheimer’s disease, respectively, this work has immediate implications for the treatment of human prion disease.

## Introduction

Prion diseases are invariably fatal neurodegenerative diseases of humans and other animals (1). The most common human prion disease, Creutzfeldt-jakob disease (CJD), accounts for ∼95% of cases; it has a sporadic etiology with a worldwide incidence of 1-2 cases per million people per year, which translates to a lifetime risk of about 1:5000. A smaller number of cases (5-10%) are due to germline mutations in *PRNP* (*Prnp* in mouse gene nomenclature), the gene encoding the prion protein (PrP^C^) (2,3). Fewer still are due to infection as a result of exposure to contaminated tissue, either by ingestion or nosocomial means (4).

The central event of prion disease is the self-templated structural rearrangement of the primarily α- helical PrP^C^ to its β-sheet enriched, disease-associated conformer, PrP^Sc^ (5,6). This three-dimensional change imparts biochemical features upon PrP^Sc^ that differentiate it from PrP^C^, including resistance to proteolytic digestion and insolubility in detergents. Following protracted and clinically silent incubation periods, accumulation of PrP^Sc^ in the central nervous system (CNS) results in the loss of neurons and eventual death of the host (1). Divergent PrP^Sc^ atomic structures with identical PrP^C^ sequences are the basis of prion strains, which cause diverse clinical and pathological outcomes by currently unknown mechanisms (5,7,8).

Extensive efforts have been devoted to identifying effective therapeutics for prion disease. Most anti-prion compounds have been discovered by exposing prion infected mouse neuroblastoma cells (ScN2a) to compound libraries and assessing changes in the levels of proteinase K (PK) resistant PrP. Molecules discovered in this manner have been shown to prolong the disease course of mice infected with mouse prions but none have yet been effective in transgenic mice expressing human PrP infected with CJD prions, or in patients (9,10). Anti-prion compounds have also been shown to have strain-specific efficacy both *in vitro* and *in vivo* (11–13), complicating efforts to develop therapeutics. It has been demonstrated that some of these molecules prevent prion propagation through a direct interaction with PrP^C^; by stabilizing its structure or by sterically occluding its interactions with PrP^Sc^ which are critical for conversion (14–16). Others are known to inhibit the conversion process by direct interaction with PrP^Sc^ (17,18). The majority of anti-prion compounds, however, do not interact with either PrP conformer and, presumably, target other molecules (19). These compounds present a more attractive pharmaceutical target as they could potentially be used to combat multiple prion strains (5,7,8).

Our laboratory recently identified phenethyl piperidines as a novel class of anti-prion compound (20). This discovery was made through application of the Drug Based Cellular Assay (DBCA) in a high throughput screen for molecules that suppress the increased sensitivity to antibiotics induced by the expression of an internally deleted form of PrP (ΔCR) (21). These compounds were refined through structure-activity relationship studies, and the most potent derivative, JZ107, can permanently cure ScN2a cells of infection by multiple prion strains. JZ107 also rescues hippocampal neurons from prion-induced synaptotoxicity *in vitro*, an important secondary assay performed on cultured hippocampal neurons (20,22,23). Phenethyl piperidine molecules do not appear to bind to PrP at concentrations used in these assays, making the identity of their interaction partners of significant interest.

In the present paper, we describe our efforts to identify the molecular target of JZ107. Through this, we discovered that ligands of the sigma receptors (σ1R and σ2R) can potently reduce the levels of PK resistant PrP, a biochemical indicator of infection, in ScN2a cells and can also prevent prion-induced retraction of dendritic spines on hippocampal neurons, a measure of synaptotoxicity. Surprisingly, however, we find that these effects are independent of direct interaction of the compounds with the sigma receptors. Because these molecules are known to cross the blood brain barrier, and some are already in clinical use for other diseases, they make excellent candidates for preclinical therapeutic studies.

## Results

### Discovery of novel anti-prion compounds

We previously described the ability of the phenethyl piperidine, JZ107, to reduce the levels of PrP^Sc^ in ScN2a cells infected with two prion strains (RML and 22L) at low micromolar concentrations (20). In an effort to identify potential non-PrP molecular targets of JZ107, we utilized the National Institute of Mental Health Psychoactive Drug Screening Program (NIMH PDSP), which assays the binding affinity of small molecules to a panel of central nervous system channels, receptors, and transporters (24). By performing radioligand binding competition assays, a list of inhibition constants (K_i_) was obtained for JZ107 and an inactive analogue, JZ103, against 37 potential target proteins (Table 1). We found that the two targets in the PDSP panel with the highest affinity for JZ107 were the sigma-1 and sigma- 2 receptors (σ_1_R and σ_2_R), which bound JZ107 with K_i_ values of 7.9 and 5.1 nM, respectively. In contrast, the K_i_ values for JZ103 binding were 339 nM (σ_1_R) and 292 nM (σ_2_R), 43x and 57x higher, respectively, than JZ107. Transcriptomic data obtained through RNAseq was used as an additional filter of this dataset, with 0.5 FPKM chosen as the lower limit for expression of each gene. Surprisingly, expression of the *Sigmar1* (σ1R) and *Tmem97* (σ2R) genes in N2a cells was found to be higher than that of any other proteins assayed in the PDSP (Table 1).

**Table 1:**
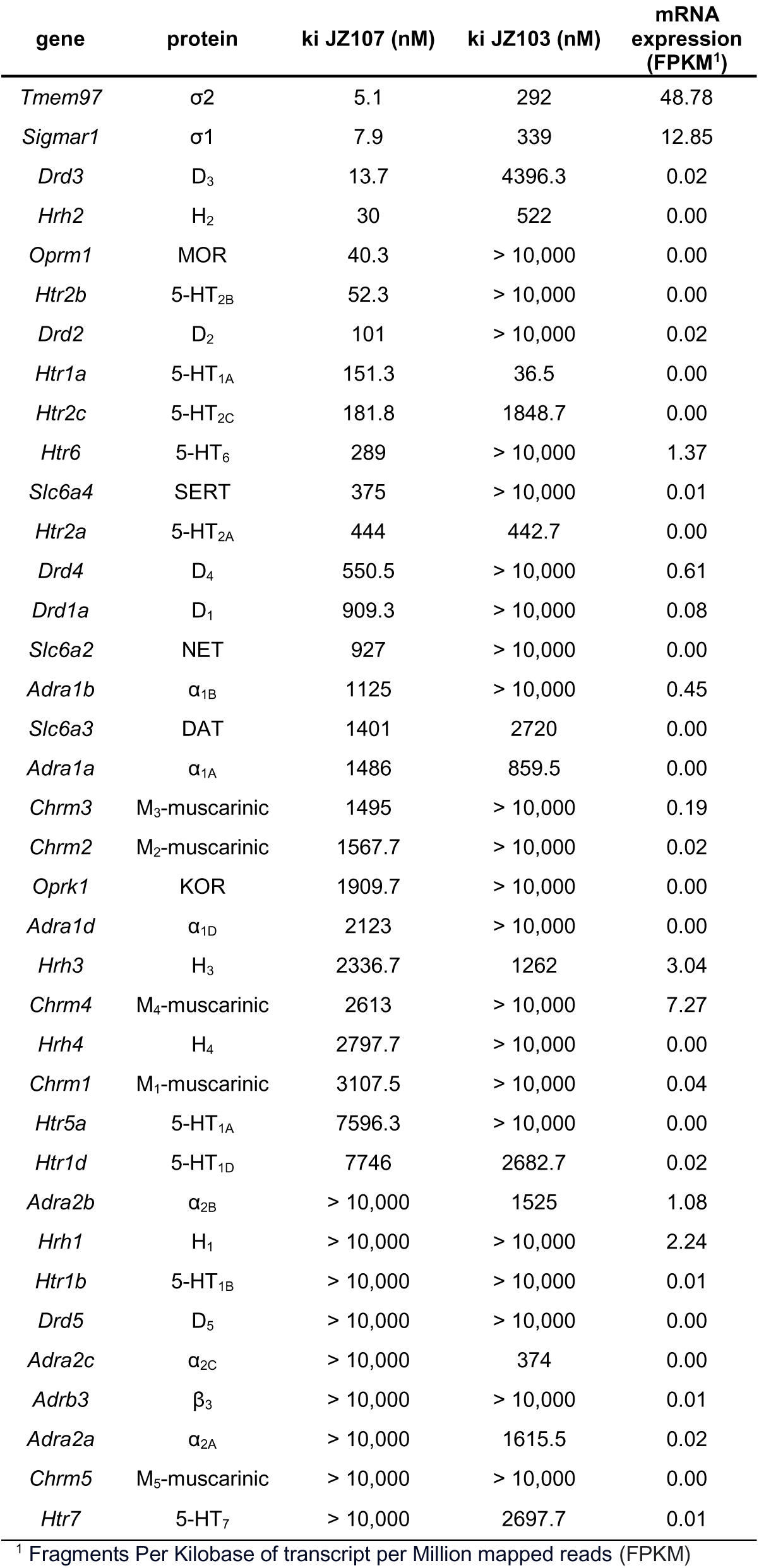
Binding affinities obtained through the PDSP and expression values in N2a cells ranked by increasing JZ107 K_i_.

| gene | protein | ki JZ107 (nM) | ki JZ103 (nM) | mRNA expression (FPKM<sup>1</sup>) |
| --- | --- | --- | --- | --- |
| *Tmem97* | $\sigma 2$ | 5.1 | 292 | 48.78 |
| *Sigmar1* | $\sigma 1$ | 7.9 | 339 | 12.85 |
| *Drd3* | D<sub>3</sub> | 13.7 | 4396.3 | 0.02 |
| *Hrh2* | H<sub>2</sub> | 30 | 522 | 0.00 |
| *Oprm1* | MOR | 40.3 | > 10,000 | 0.00 |
| *Htr2b* | 5-HT<sub>2B</sub> | 52.3 | > 10,000 | 0.00 |
| *Drd2* | D<sub>2</sub> | 101 | > 10,000 | 0.02 |
| *Htr1a* | 5-HT<sub>1A</sub> | 151.3 | 36.5 | 0.00 |
| *Htr2c* | 5-HT<sub>2C</sub> | 181.8 | 1848.7 | 0.00 |
| *Htr6* | 5-HT<sub>6</sub> | 289 | > 10,000 | 1.37 |
| *Slc6a4* | SERT | 375 | > 10,000 | 0.01 |
| *Htr2a* | 5-HT<sub>2A</sub> | 444 | 442.7 | 0.00 |
| *Drd4* | D<sub>4</sub> | 550.5 | > 10,000 | 0.61 |
| *Drd1a* | D<sub>1</sub> | 909.3 | > 10,000 | 0.08 |
| *Slc6a2* | NET | 927 | > 10,000 | 0.00 |
| *Adra1b* | $\alpha_{1B}$ | 1125 | > 10,000 | 0.45 |
| *Slc6a3* | DAT | 1401 | 2720 | 0.00 |
| *Adra1a* | $\alpha_{1A}$ | 1486 | 859.5 | 0.00 |
| *Chrm3* | M<sub>3</sub>-muscarinic | 1495 | > 10,000 | 0.19 |
| *Chrm2* | M<sub>2</sub>-muscarinic | 1567.7 | > 10,000 | 0.02 |
| *Oprk1* | KOR | 1909.7 | > 10,000 | 0.00 |
| *Adra1d* | $\alpha_{1D}$ | 2123 | > 10,000 | 0.00 |
| *Hrh3* | H<sub>3</sub> | 2336.7 | 1262 | 3.04 |
| *Chrm4* | M<sub>4</sub>-muscarinic | 2613 | > 10,000 | 7.27 |
| *Hrh4* | H<sub>4</sub> | 2797.7 | > 10,000 | 0.00 |
| *Chrm1* | M<sub>1</sub>-muscarinic | 3107.5 | > 10,000 | 0.04 |
| *Htr5a* | 5-HT<sub>1A</sub> | 7596.3 | > 10,000 | 0.00 |
| *Htr1d* | 5-HT<sub>1D</sub> | 7746 | 2682.7 | 0.02 |
| *Adra2b* | $\alpha_{2B}$ | > 10,000 | 1525 | 1.08 |
| *Hrh1* | H<sub>1</sub> | > 10,000 | > 10,000 | 2.24 |
| *Htr1b* | 5-HT<sub>1B</sub> | > 10,000 | > 10,000 | 0.01 |
| *Drd5* | D<sub>5</sub> | > 10,000 | > 10,000 | 0.00 |
| *Adra2c* | $\alpha_{2C}$ | > 10,000 | 374 | 0.00 |
| *Adrb3* | $\beta_3$ | > 10,000 | > 10,000 | 0.01 |
| *Adra2a* | $\alpha_{2A}$ | > 10,000 | 1615.5 | 0.02 |
| *Chrm5* | M<sub>5</sub>-muscarinic | > 10,000 | > 10,000 | 0.00 |
| *Htr7* | 5-HT<sub>7</sub> | > 10,000 | 2697.7 | 0.01 |

<sup>1</sup> Fragments Per Kilobase of transcript per Million mapped reads (FPKM)

These results led us to test several known sigma receptor ligands for their ability to reduce the levels of PK resistant PrP. RML infected ScN2a cells were cultured in the presence of increasing concentrations of each compound or DMSO vehicle for a total of seven days, with passaging on the third day. Cell lysates were then exposed to 10 µg/ml of PK before western blotting while MTT assays were performed in parallel as a measure of cell viability. Using this approach, we identified ten novel anti-prion compounds: PD 144418 (25) (EC_50_ = 5.2 µM) (Figure 1A); BD1047 (26) (EC_50_ = 19 µM) (Figure 1B); BD1063 (26) (EC_50_ = 12.3 µM) (Figure 1C); PB-28 (27) (EC_50_ = 5.1 µM) (Figure 1D); rimcazole (BW234U) (28) (EC_50_ = 3.5 µM) (Figure 1E); haloperidol (29) (EC_50_ = 13.2 µM) (Figure 1F); SA4503 (Cutamesine) (30) (EC_50_ = 27.2 µM) (Figure 1G), ANAVEX2-73 (Blarcamesine) (31) (EC_50_ = 39.5 µM) (Figure 1H), (+)-pentazocine (32) (EC_50_ = 35.8 µM) (Figure 1I), and ditolylguanidine (DTG) (33) (EC_50_ = 68.1 µM) (Figure 1J). BMY-14802 (34) was found to be ineffective in lowering the levels of PK resistant PrP at concentrations up to 50 µM, and PrP^Sc^ reduction at higher concentrations paralleled a loss of cell viability (Figure 1K). For ease of visual comparison, dose response curves are plotted together (Figure 1L), and all data, including compound structures, are compiled in Table 2 along with corresponding data for JZ107 and JZ013 (20). Based on MTT assays of cellular toxicity (Figure 1, dashed lines), molecules with anti-prion activity exhibited LC_50_ values ranging from 7.8 - >50 µM, with corresponding therapeutic indices (TI = LC_50_/EC_50_) of 1.5 - >4.2 (Table 2).

**Figure 1:**
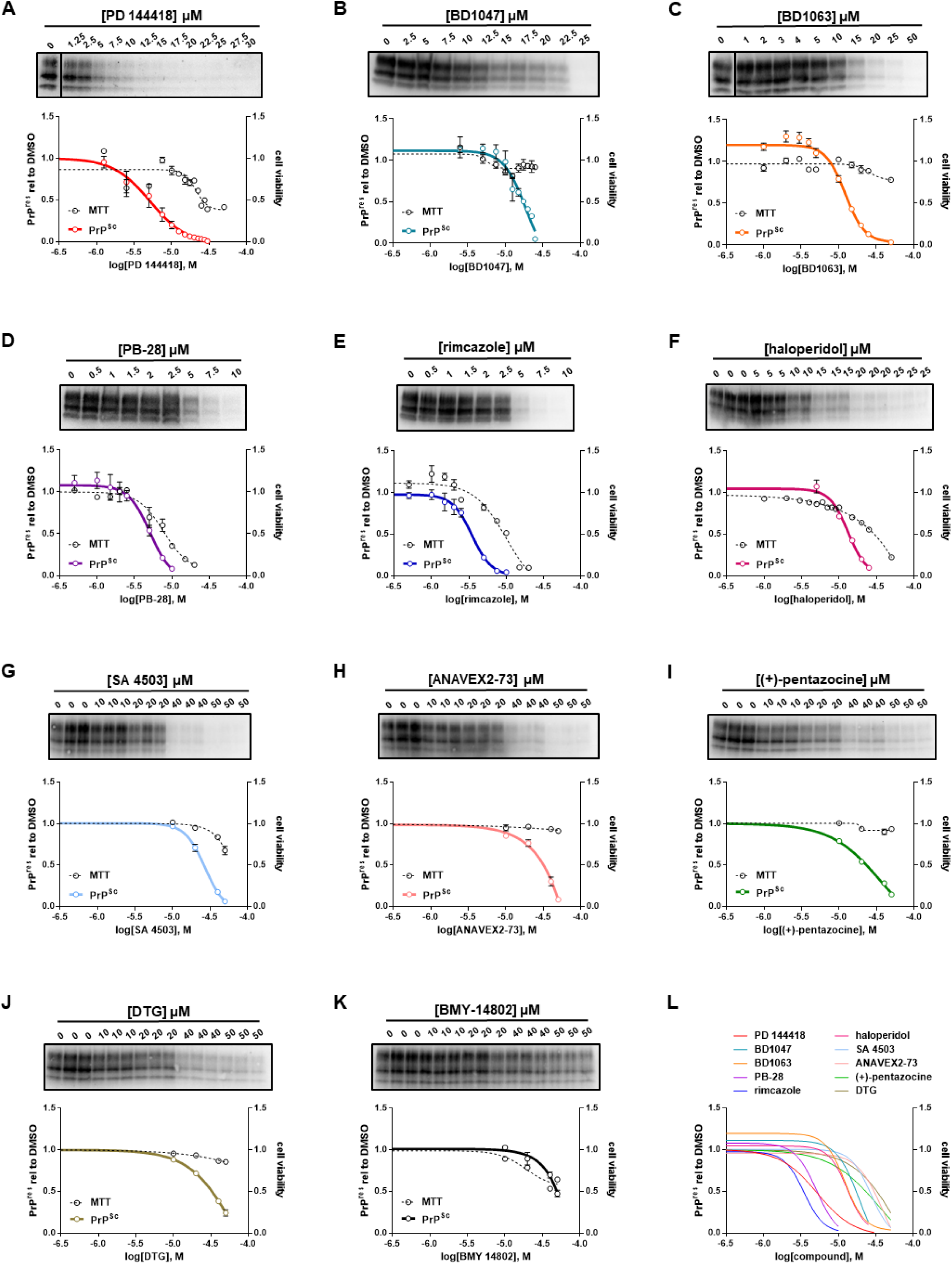
Sigma receptor ligands reduce the levels of PK resistant PrP in ScN2a-RML cells. N2a cells chronically infected with RML prions were incubated with the indicated concentrations of compound for a total of seven days, before being lysed for PK digestion and analysis by western blot. Cultures treated in parallel were subjected to MTT assay. All data points represent three independent replicates. **A)** PD 144418; **B)** BD1047; **C)** BD1063; **D)** PB-28; **E)** rimcazole; **F)** haloperidol; **G)** SA 4503; **H)** ANAVEX2-73; **I)** (+)-pentazocine; **J)** DTG; **K)** BMY 14802; **L)** EC_50_ curves of A-K plotted together. All curves were fit by least squares regression using GraphPad software, n = 3 replicates.

**Table 2:**
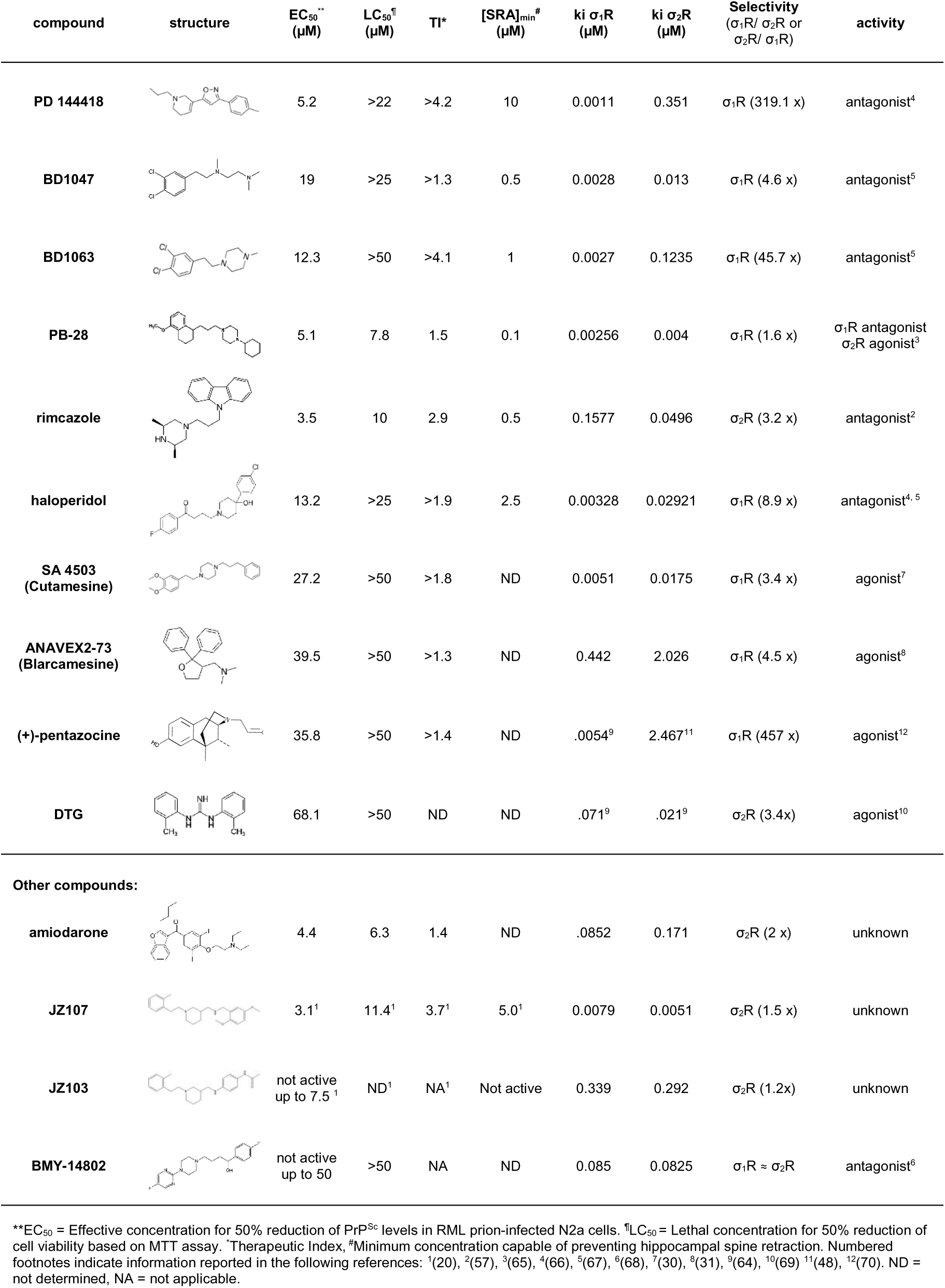
Structures and pharmacological properties of compounds used in this study.

### Sigma receptor binding characteristics of anti-prion compounds

K_i_ values reported in the literature for sigma receptor ligands are often difficult to compare, as they have been obtained using different assay methods and/or source tissues. We therefore submitted our set of anti-prion compounds to the PDSP to experimentally verify their sigma receptor binding characteristics. The PDSP assays the ability of test compounds to compete with binding of [^3^H]-(+)- pentazocine and [^3^H]-DTG to membrane preparations of receptor-expressing HEK293T cells to determine K_i_ values for σ_1_R and σ_2_R, respectively. We observed that, irrespective of anti-prion activity, all compounds bound to both σ_1_R and σ_2_R; most with sub-micromolar affinity (Table 2). Of the ligands with anti-prion activity, PB-28 and JZ107 had similar affinities for σ_1_R and σ_2_R (1.6X and 1.5X selectivity for σ_1_R and σ_2_R, respectively). Showing slight selectivity for σ_1_R were BD1047 (4.6x), haloperidol (8.9x), SA 4503 (3.4x), and ANAVEX2-73 (4.5x), while rimcazole and DTG displayed the opposite preference (3.2X and 3.4X for σ_2_R, respectively). Finally, three ligands were highly selective for σ_1_R: PD 144418 (319.1x), BD1063 (45.7x), and (+)-pentazocine (457x).

We attempted to correlate the observed anti-prion effects with the categorization of these compounds as agonists or antagonists of the sigma receptors. This categorization must be regarded cautiously, since there is no agreed upon biochemical or physiological activity of either receptor to serve as a criterion for assigning the pharmacological effects of its ligands (Table 2). Although most compounds with anti-prion activity have been classified as antagonists in published literature, some (SA 4503 and ANAVEX2-73) have been classified as agonists, and one (PB28) as an antagonist of σ_1_R and an agonist of σ_2_R. We also assessed the relationship between the observed anti-prion activities of these molecules with their binding affinity for the two sigma receptors. When we plotted the EC_50_ values determined using ScN2a cells against K_i_ values for σ_1_R and σ_2_R determined using the PDSP, we observed no correlation between these two parameters for either receptor (Figure 2). Taken together, these results fail to confirm a correlation between the anti-prion potency of the ten ligands and their affinity for or purported pharmacological action on σ_1_R and σ_2_R.

**Figure 2:**
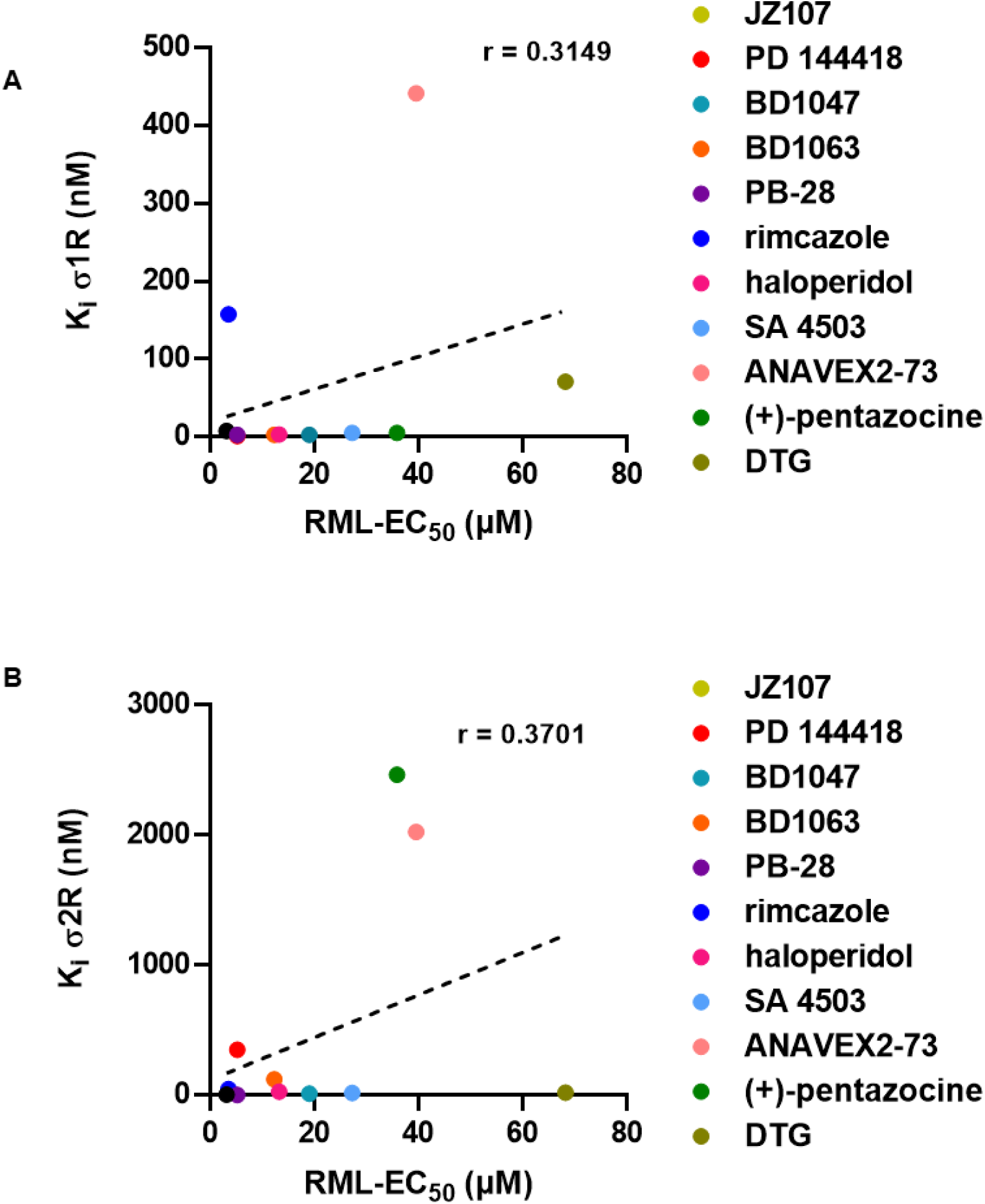
Correlation of sigma receptor binding affinity and anti-prion activity. The corresponding K_i_ for each compound for σ1R or σ2R as determined by the PDSP is plotted against the EC_50_ determined against RML prions. PDSP derived K_i_ values of (+)-pentazocine and DTG were determined by another group (48,64). **A)** σ1R K_i_ vs RML EC_50_: r = 0.3149, p = 0.3455. **B)** σ2R K_i_ vs RML EC_50_: r = 0.3701, p = 0.2626. Correlation analysis was performed using GraphPad software.

### Sigma receptor knockout does not abrogate anti-prion effects

To investigate the role of σ_1_R and σ_2_R in prion propagation and in the anti-prion effects of their ligands, we edited both the *Sigmar1* and *Tmem97* genes in ScN2a cells using CRISPR/Cas9 to disrupt the respective open reading frames. Since N2a cells are well known to display clonal variations in prion infection susceptibility and propagation (35), we utilized a CRISPR/Cas9 platform capable of high-efficiency gene editing without the need for cloning (see Experimental Procedures). As a control, we used a sgRNA targeting the Rosa26 safe harbor locus. Editing efficiency was determined by Sanger sequencing of PCR amplicons and deconvolution using the Inference of CRISPR Edits (ICE) tool (36). The editing efficiency (percentage of open reading frames with indels), and the knock-out score (percentage of edits producing a frameshift) was > 94% for both *Sigmar1* and *Tmem97* (Figure 3A). Quantitative RT-PCR analysis demonstrated that *Sigmar1* and *Tmem97* transcripts underwent incomplete nonsense mediated decay, with transcript levels of *Sigmar1* and *Tmem97* decreasing relative to Rosa26 targeted cells (Figure 3B). Confirming nucleotide analysis, σ_1_R and σ_2_R could not be detected in *Sigmar1* + *Tmem97* knockout cells by western blot (Figure 3C). These genetic manipulations had no effect on either the transcript level of *Prnp*, (Figure 3B), or upon the level of total or PK resistant PrP (Figure 3C, D). These results indicate that genetic reduction of σ_1_R and σ_2_R expression in ScN2a cells has no significant effect on the basal levels of PrP^Sc^ in the absence of sigma receptor ligands.

**Figure 3:**
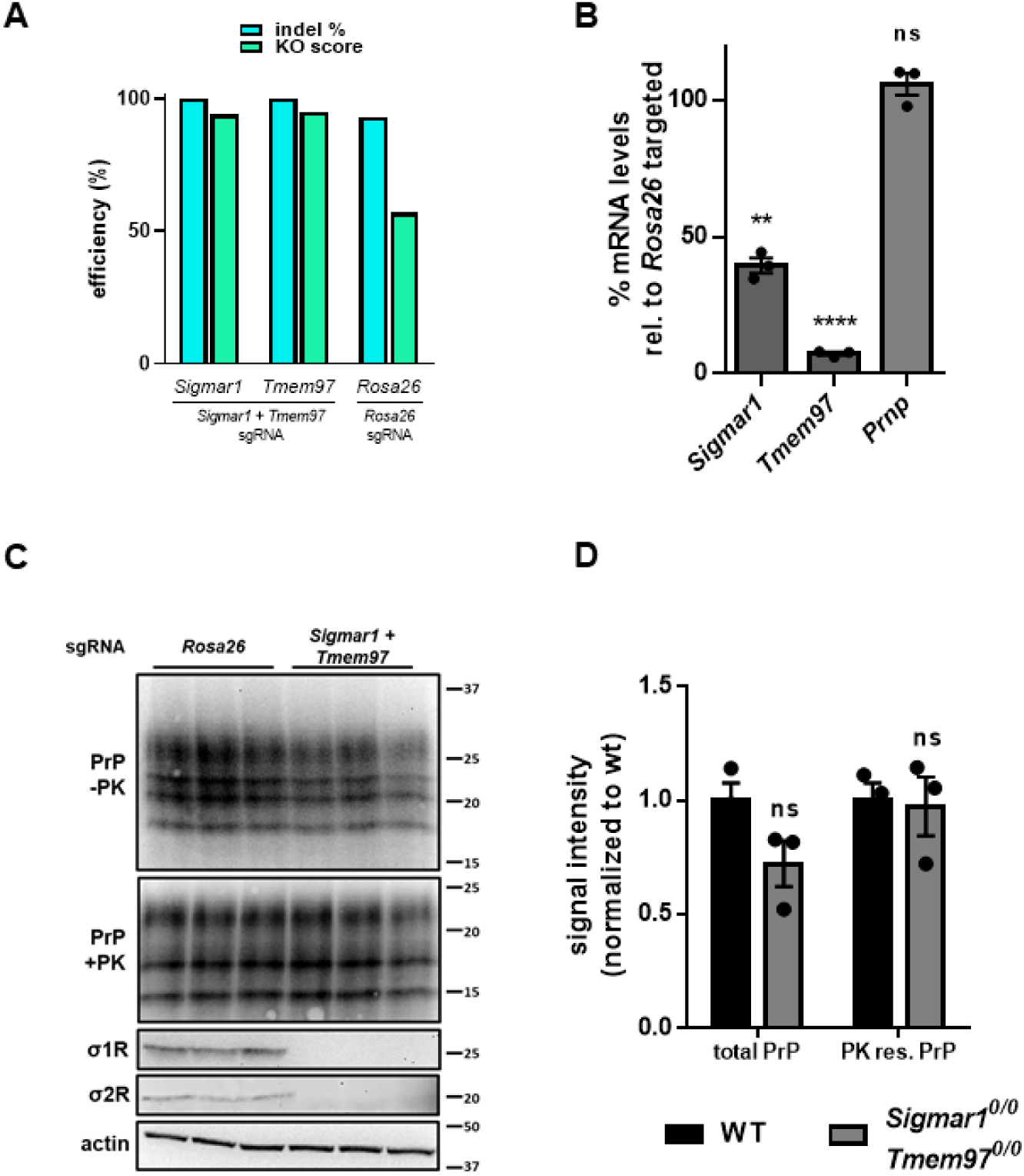
Combined CRISPR/Cas9 mediated editing of the σ1R and σ2R genes has no effect on basal levels of PrP^C^ or PrP^Sc^. **>A)** Inference of CRISPR Edits (ICE) analysis of *Sigmar1* and *Tmem97* gene disruption. Following PCR amplification and cleanup of targeted loci, ICE analysis allows for the deconvolution of Sanger sequencing data to provide indel % (editing efficiency) and a knockout score (KO: proportion of edits that result in a frame shift). The Rosa26 safe harbor locus is targeted with a single guide RNA as a control. **B)** Quantitative RT-PCR analysis using the ΔΔC_t_ method. **C)** High knockout efficiency is demonstrated by undetectable levels of σ1R and σ2R protein in double targeted cells by western blot. Blotting for PrP with and without PK demonstrates that levels are unaffected by the ablation of *Sigmar1* and *Tmem97*. **D)** Quantification of PrP blots in **C**. For all experiments n = 3 replicates. p < 0.0001 = ****; p < 0.001 = ***; p < 0.01 = **; p < 0.05 = *; ns = not significant using a two-tailed students t-test.

We next sought to determine if knockout of σ_1_R and σ_2_R influenced the ability of sigma receptor ligands to reduce PrP^Sc^ levels. For these experiments, we subjected these cells to our standard drug treatment regimen using the determined EC_50_ of each compound against RML prions in unedited cells (Figure 1; Table 2). Surprisingly, we found that dual knockout of *Sigmar1* and *Tmem97* did not affect the ability of any of the compounds to reduce the levels of PK resistant PrP (Figure 4). These data demonstrate that, despite being sigma receptor ligands, the anti-prion activity of these compounds does not require expression of σ_1_R or σ_2_R.

**Figure 4:**
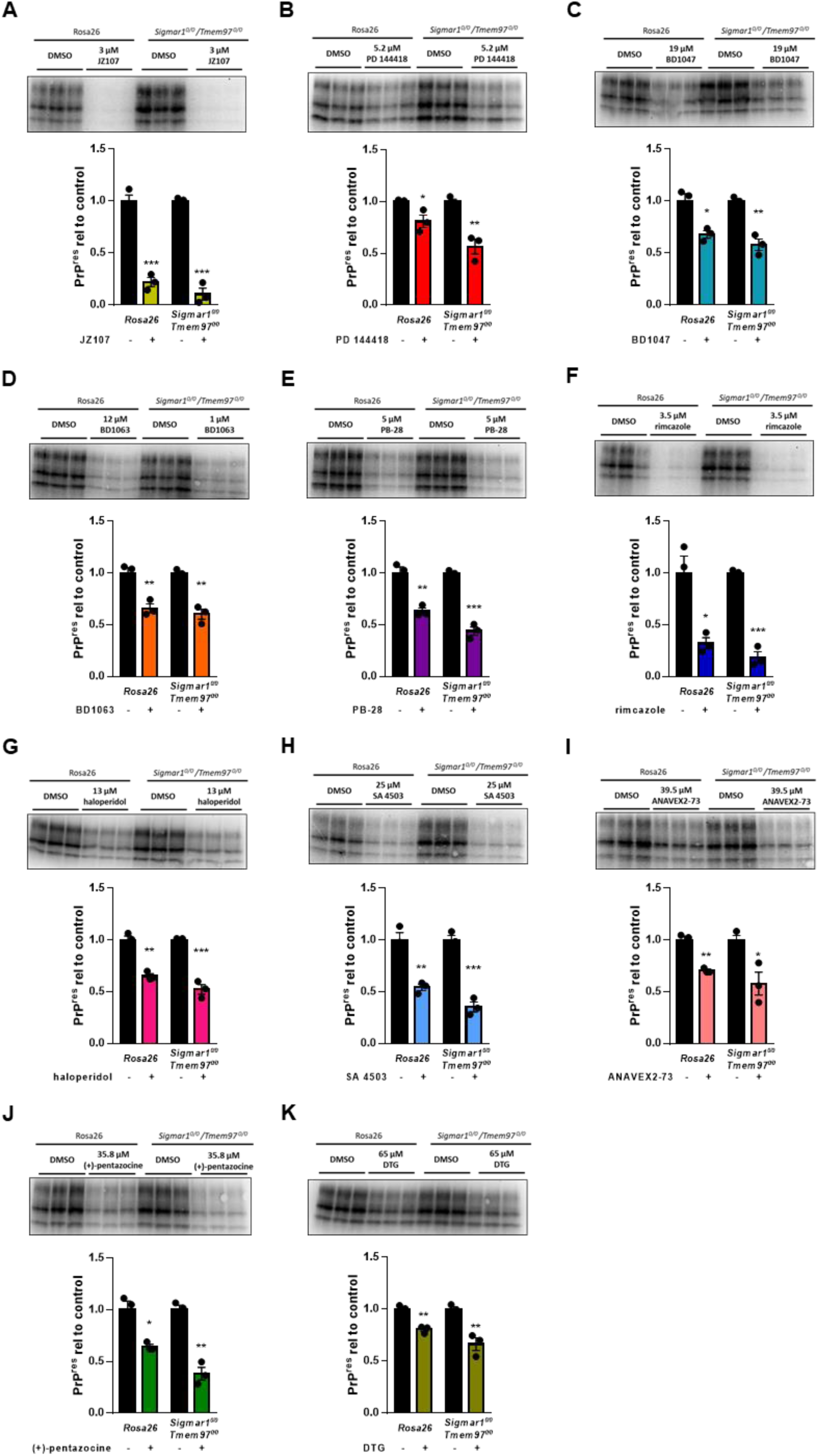
Elimination of σ1R and σ2R expression does not alter the anti-prion effects of sigma receptor ligands. Compound efficacy against RML prions in chronically infected *Sigmar1* + *Tmem97* KO cells. For each compound, the EC_50_ determined against RML prions in N2a cells was used. Levels of PK resistant PrP relative to DMSO treated controls are plotted (mean ± SEM). **A)** JZ107; **B)** PD 144418; **C)** BD1047; **D)** BD1063; **E)** PB-28; **F)** rimcazole; **G)** haloperidol; **H)** SA 4503; **I)** ANAVEX2-73; **J)** (+)- pentazocine; **K)** DTG. For all experiments n = 3 replicates. p < 0.0001 = ****; p < 0.001 = ***; p < 0.01= **; p < 0.05 = *; ns = not significant using the Holm-Sidak test for multiple comparisons.

### Other binding partners are not responsible for anti-prion effects

While σ_1_R or σ_2_R were the targets identified by the PDSP that were most highly expressed in N2a cells and that had the highest affinity for the test compound JZ107, (Table 1), other proteins that interact with one or more of sigma receptor ligands are expressed at lower levels (Figure 5A; Table 1; Supplementary Table 1). Using the same strategy employed for *Sigmar1* and *Tmem97*, we disrupted the open reading frames of *Htr6*, *Drd4*, *Hrh1*, *Hrh3*, and *Chrm4*. Attempts to detect the corresponding proteins by western blot were unsuccessful, likely due to their low levels of expression in N2a cells (Table 1). However, disruption of the open reading frames observed through ICE analysis gives us confidence that these genes are no longer functional (Figure 5B). Again using each compound at its previously determined EC_50_, we were unable to demonstrate the requirement for any of these proteins for the observed anti-prion effects of the tested compounds (Figure 5C-N).

**Figure 5:**
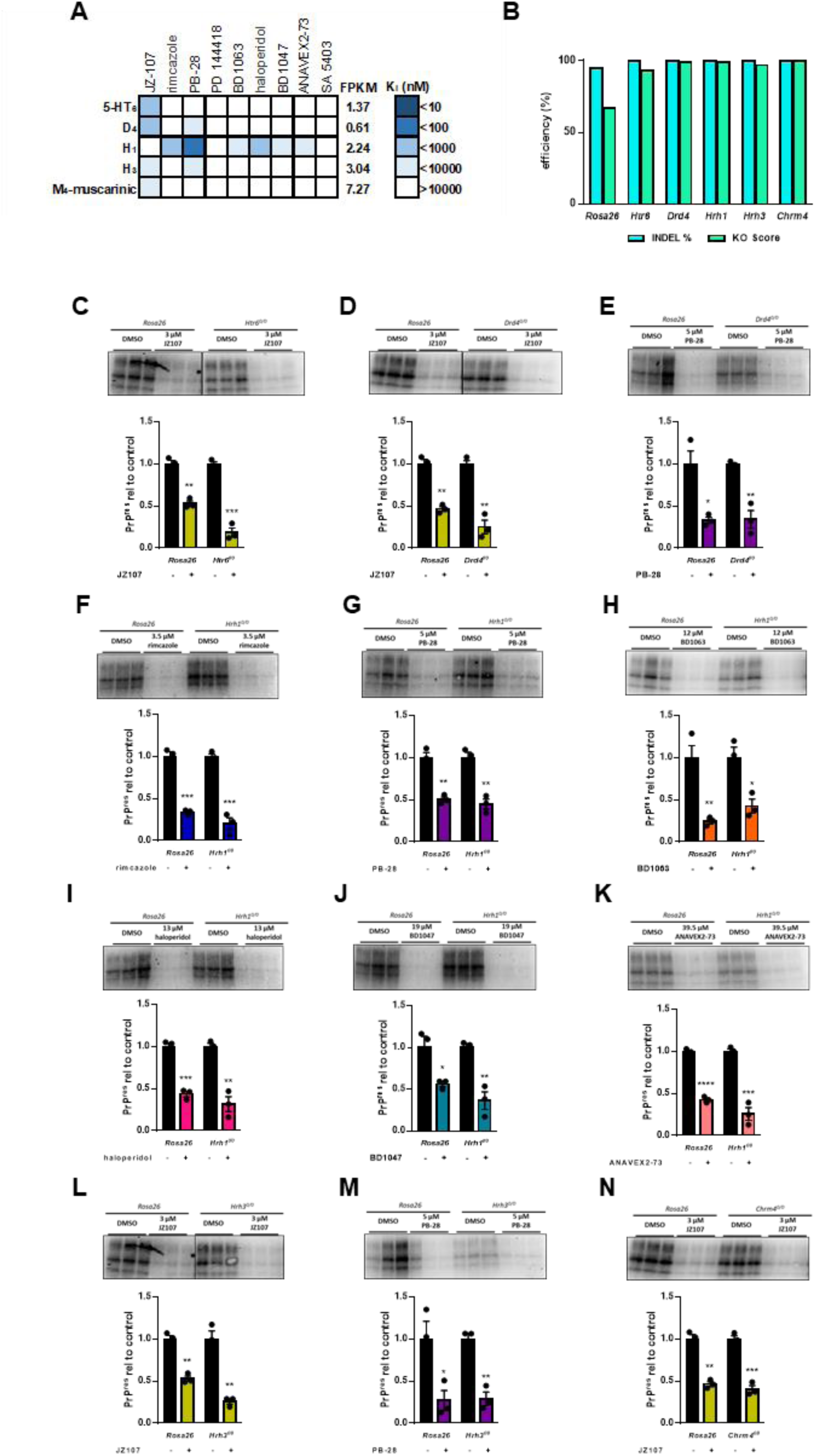
Other receptors identified in the PDSP do not mediate the observed anti-prion effects. **A)** Graphical representation of active compound K_i_ values for other receptors known to be expressed in N2a cells (FPKM > 0.5). **B)** Inference of CRISPR Edits (ICE) analysis of gene disruption. The Rosa26 safe harbor locus is targeted with a single guide RNA as a control. **C-N)** Compound efficacy against RML prions in chronically infected KO cells. For each compound, the EC_50_ determined against RML prions in N2a cells was used. Levels of PK resistant PrP are expressed relative to DMSO treated controls in Rosa26 targeted cells vs KO cells ± SEM : **C)** JZ107 against *Htr6^0/0^* cells; **D)** JZ107 against *Drd4^0/0^* cells; **E)** PB-28 against *Drd4^0/0^* cells; **F)** rimcazole against *Hrh1^0/0^*cells; **G)** PB-28 against *Hrh1^0/0^* cells; **H)** BD1063 against *Hrh1^0/0^* cells; **I)** haloperidol against *Hrh1^0/0^* cells; **J)** BD1047 against *Hrh1^0/0^* cells; **K)** ANAVEX2-73 against *Hrh1^0/0^* cells; **L)** JZ107 against *Hrh3^0/0^* cells; **M)** PB-28 against *Hrh3^0/0^* cells; **N)** JZ107 against *Chrm4^0/0^* cells. Assays shown in **C** and **L**, and in **D** and **N** were done using the same western blot, necessitating the use of the same Rosa26 control images. For all experiments n = 3 replicates. p < 0.0001 = ****; p < 0.001 = ***; p < 0.01 = **; p < 0.05 = *; ns = not significant using the Holm-Sidak test for multiple comparisons.

### Phospholipidosis as a potential anti-prion mechanism

σ_1_R and σ_2_R were recently identified in a screen for host molecules that interact with SARS-CoV-2 proteins and it was demonstrated that known sigma receptor ligands, including some under investigation here, possessed antiviral activity (37). However, a subsequent study found that these effects were not correlated with binding affinity for σ_1_R or σ_2_R, but instead depended on the ability of these molecules to induce phospholipidosis (38). A phenomenon that often confounds drug discovery efforts, phospholipidosis occurs due to the capacity of cationic, amphiphilic molecules to accumulate in endosomes and lysosomes, resulting in disruptions of lipid processing and alterations in the morphology of these compartments (39). Because lysosomal function is known to be important for PrP^Sc^ degradation, and some of the molecules under investigation here are reported inducers of phospholipidosis, we next explored this as a potential mechanism of action of these anti-prion compounds (38,40).

Interestingly, in addition to being a potent inducer of phospholipidosis (38), the antiarrhythmic medication amiodarone has been previously identified as an anti-prion compound (41) and sigma receptor ligand (42). In our hands, amiodarone has an EC_50_ of 4.4 µM and a LC_50_ of 6.3 µM (Figure 6A), and K_i_ values of 85.3 nM and 171 nM for σ_1_R and σ_2_R, respectively (Table 2). Here, we used amiodarone as a positive control in phospholipidosis assays, which were monitored using LipidTox, a fluorescent reagent that stains intracellular lipid accumulations (43). Similar results were obtained with an alternative reporter, NBD-PE (not shown, (38)). Of the most potent compounds, only rimcazole induced phospholipidosis at levels > 50% that of amiodarone after incubation with N2a cells at 10 µM for 24 hours (Figure 6B, C). Further, the level of phospholipidosis induced by these compounds did not correlate with their anti-prion activity in ScN2a cells (r = −0.4436, p = 0.2709) (Figure 6D). These results indicate that phospholipidosis itself is unlikely to be a major mechanism underlying the anti-prion effects of the sigma receptor ligands analyzed here, although we cannot rule out that other alterations in lysosomal or autophagosomal pathways could be a contributing factor to the action of some of the compounds.

**Figure 6:**
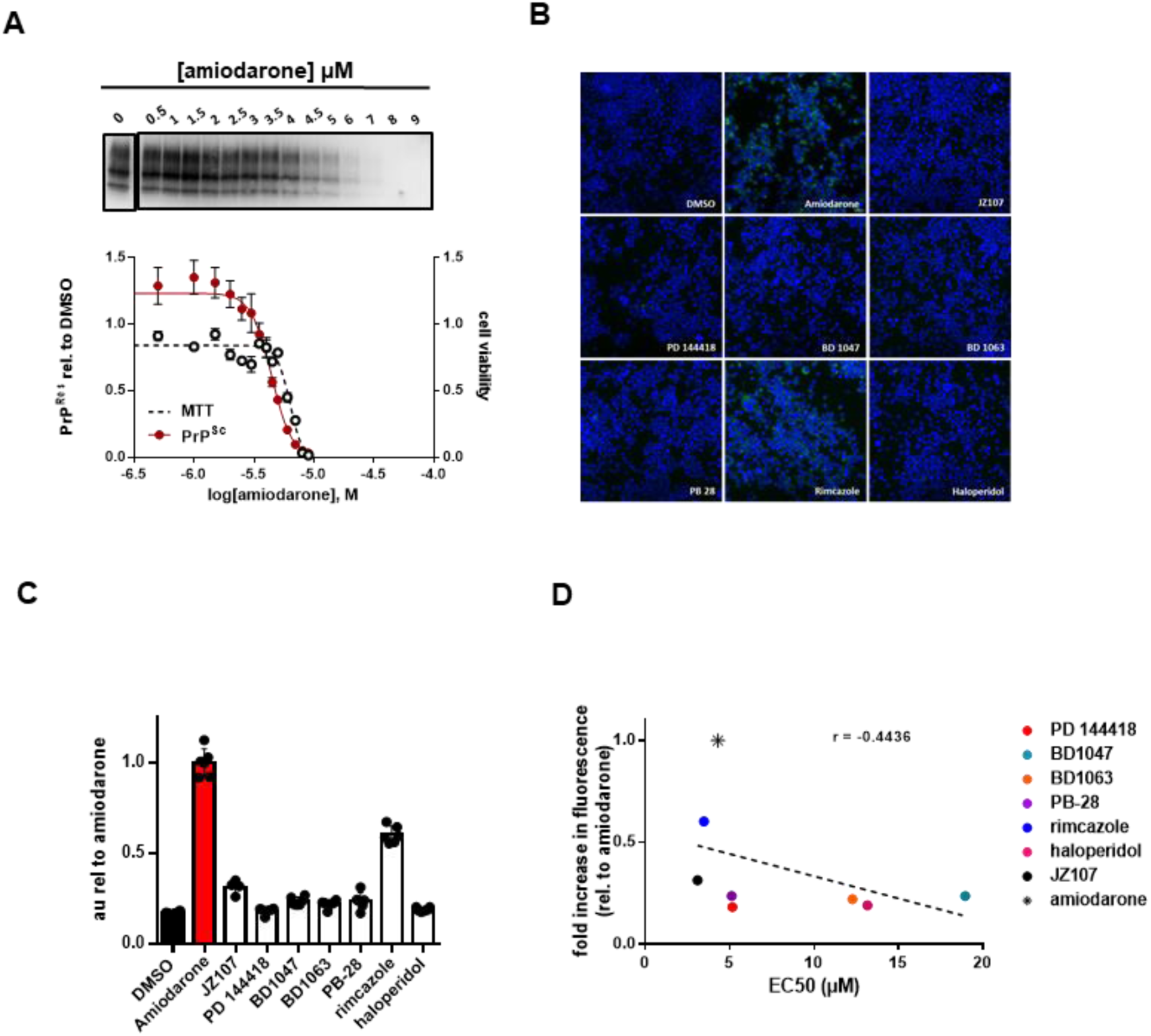
Phospholipidosis induction is not correlated with anti-prion activity. **A)** N2a cells chronically infected with RML prions were incubated with the indicated concentrations of amiodarone for a total of seven days, before being lysed for PK digestion and analysis by western blot. Cultures treated in parallel were subjected to MTT assay. All data points represent three independent replicates. **B)** Uninfected N2a cells were incubated in 10 µM of the indicated compound with LipidTox for 24 hours. Green fluorescence represents LipidTox staining, blue represents Hoechst nuclei staining. **C)** Total LipidTox fluorescence was normalized to signal derived from Hoechest. **D)** For each compound, the level of phospholipidosis induction relative to DMSO is plotted against the EC_50_ determined against RML prions. r = −0.4436, p = 0.2709. Correlation analysis was performed using GraphPad software.

### Sigma receptor ligands do not alter the levels or sub-cellular localization of PrP^C^

To further probe the mechanism of action of the sigma receptor ligands, we tested their effects on the total levels and sub-cellular localization of PrP^C^, factors that could potentially influence cellular content of PrP^Sc^. Focusing on compounds with an EC_50_ of < 20 µM, we found that none significantly altered the total levels of PrP^C^ assessed by western blotting when tested at their EC_50_ values for anti-prion activity (Figure 7A; Supplementary Figure 1). The compounds also had no observable effect on the sub-cellular localization of PrP^C^ based on immunofluorescence staining (Figure 7B).

**Figure 7:**
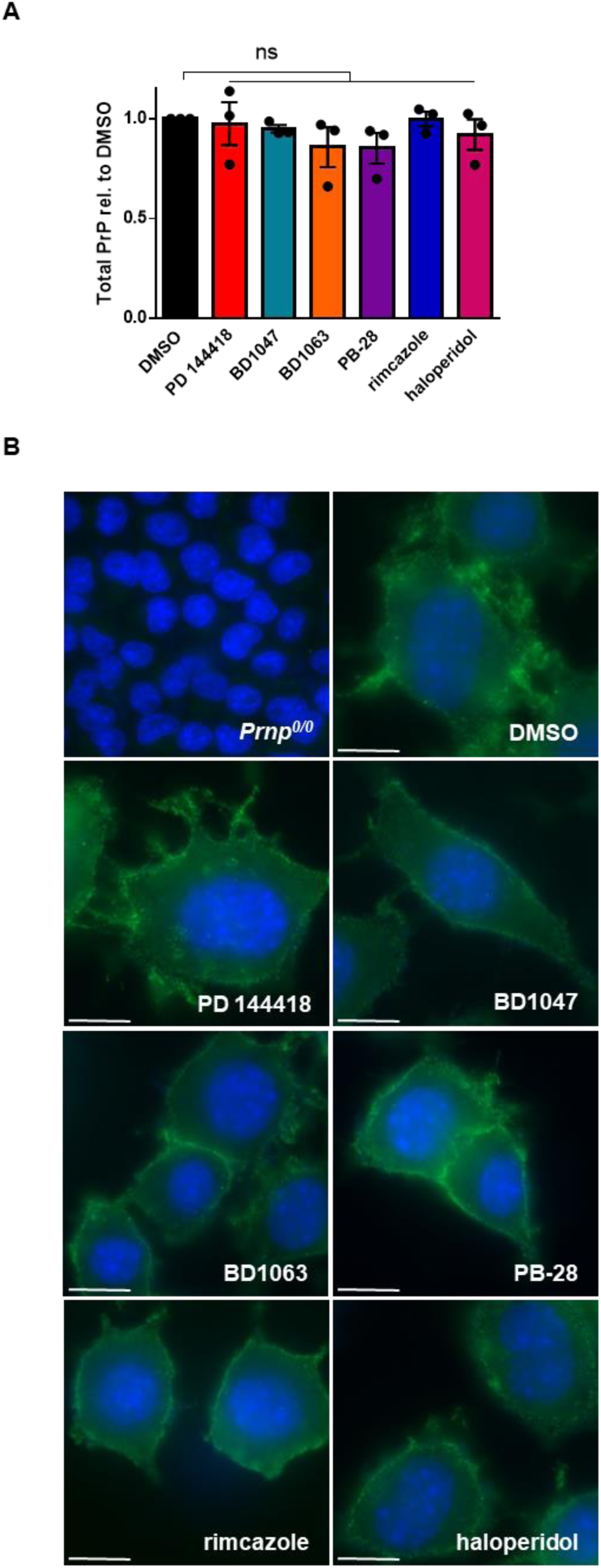
Anti-prion compounds do not alter PrP^C^ levels or sub-cellular localization. **A)** Effect of compounds on total PrP levels in uninfected N2a cells. For each compound, the EC_50_ determined against RML prions in Figure 2 was used. Following incubation with each compound for seven days, levels of total PrP, compared to DMSO treatment, were plotted as mean ± SEM. **B)** Compounds do not alter the cell-surface localization of PrP^C^. For each compound, the EC_50_ determined against RML prions was used. PrP_C_ was imaged by immunofluorescence staining using D18. DAPI is used as a nuclear counter stain. Scale bar = 10 μm. For all experiments, n = 3 replicates. p < 0.0001 = ****; p < 0.001 = ***; p < 0.01 = **; p < 0.05 = *; ns = not significant using a two-tailed students t-test.

### Prion strain specificity of sigma receptor ligands

The efficacy of some anti-prion compounds has been shown to be strain dependent (11,44). To examine the potential for strain specificity, we used N2a cells chronically infected with a second strain of prions (22L), again focusing on compounds that displayed an EC_50_ of < 20 µM against RML prions. We observed that three of these compounds have efficacies against 22L prions that are comparable to those against RML prions: When tested at their EC_50_ values for RML prions, BD1063 reduced PK resistant PrP to 63.3 ± 1.2% (p = < 0.0001), PB-28 by 52.6 ± 2.6% (p = < 0.0001), and haloperidol by 43.0 ± 8.6% (p = 0.0007) of that observed in vehicle treated cells (Figure 8A, Supplementary Figure 2C, D, F). Two molecules were more effective against 22L prions than RML prions, resulting in reductions in PK resistant PrP by BD1047 to 19.1 ± 10.5% (p = 0.0004) and by rimcazole to 13.5 ± 7.8% (p = < 0.0001) of that seen in controls (Figure 8A, Supplementary Figure 2B, E). PD 144418, however, was ineffective against 22L prions at the EC_50_ determined against RML prions (Figure 8A; Supplementary Figure 2A). However, a dose response curve using 22L infected ScN2a cells revealed that PD 144418 is effective at higher concentrations, with an EC_50_ of 12.6 µM against this strain compared to an EC_50_ of 5.2 µM against RML prions (Figure 8B).

**Figure 8:**
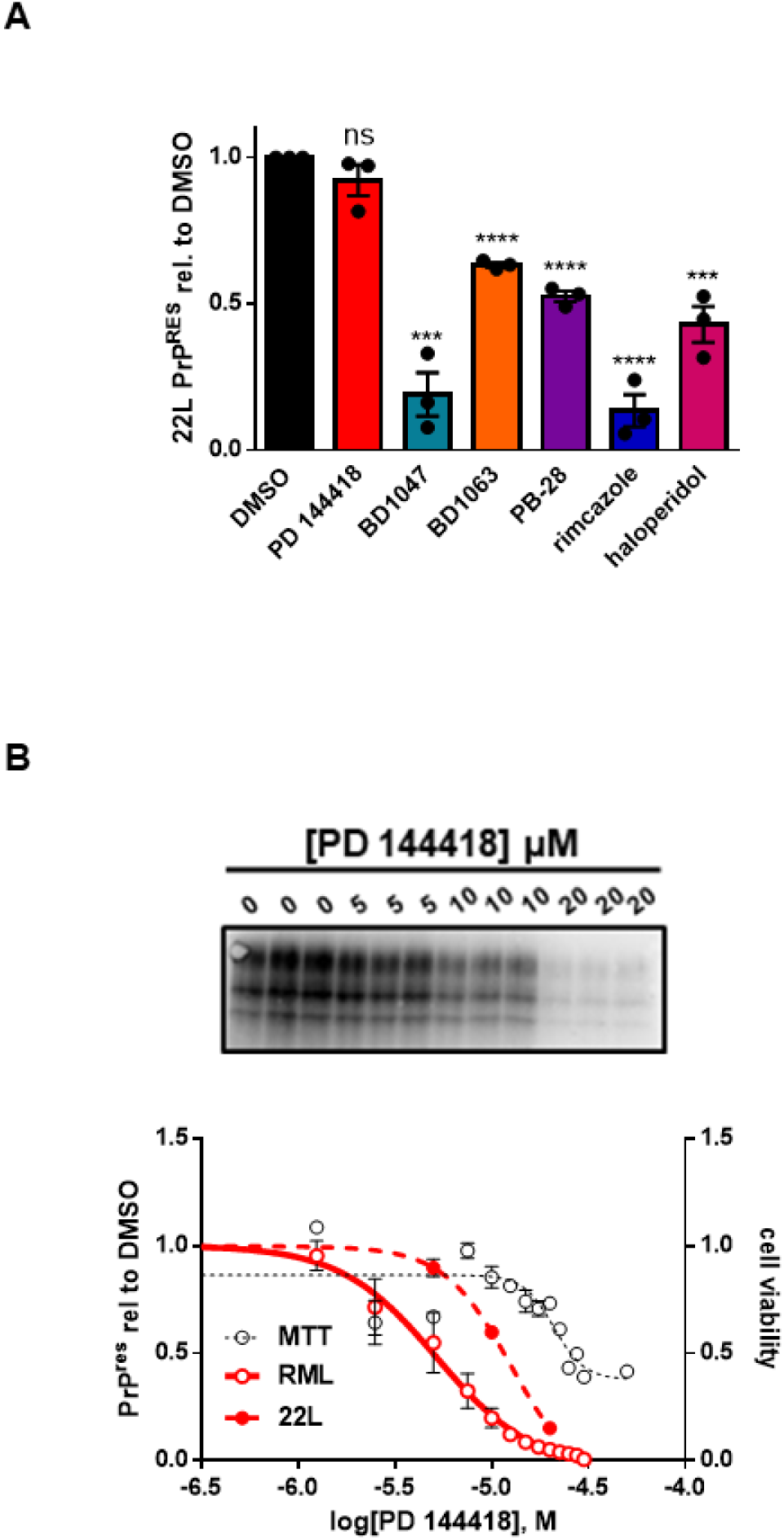
Anti-prion compounds do not exhibit strain specificity. **A)** Compound efficacy against 22L prions in chronically infected N2a cells. For each compound, the EC_50_ determined against RML prions was used. Levels of PK resistant PrP, compared to DMSO treatment, were plotted as mean ± SEM. **B)** Dose response curves of PK resistant PrP remaining after incubation with increasing concentrations of PD 144418. RML and MTT values are replotted from Figure 1A. For all experiments, n = 3 replicates. p < 0.0001 = ****; p < 0.001 = ***; p < 0.01 = **; p < 0.05 = *; ns = not significant using a two-tailed students t-test.

### Sigma receptor ligands prevent prion-induced synaptotoxicity

We previously demonstrated that exposure of cultured hippocampal neurons to prion infected brain homogenates or purified PrP^Sc^ causes a rapid (<12 hours) and dramatic retraction of dendritic spines (22,45,46). This phenomenon, which we have used as an assay of the synaptotoxic activity of prions, is dependent on the presence of cell-surface PrP^C^ and is blocked by NMDA receptor antagonists and p38 MAPK inhibitors. Our results suggest that rapid conversion of PrP^C^ to PrP^Sc^ on the cell surface triggers an intracellular signaling cascade that ultimately leads to collapse of the actin cytoskeleton within dendritic spines. In this scenario, compounds that inhibit the conversion of PrP^C^ to PrP^Sc^ would be predicted to prevent spine collapse upon prion exposure. To test this prediction, hippocampal neurons (21 DIV) were pre-incubated with one of the six most potent compounds or DMSO vehicle for 2 hours before treatment with either purified RML prions or mock-purified material from uninfected, age-matched control brains (Supplementary Figure 3). Following 24 hours of prion exposure in the continued presence of the test compound, spine density was determined by immunofluorescence staining for F-actin. As expected, the application of purified prions to these cultures in the absence of test compound caused dendritic spine collapse, reducing spine density to less than 40% of that observed in cultures treated with mock-purified material (Figure 9; mock vs prion). Strikingly, all six sigma receptor ligands tested in this assay were able to significantly prevent spine loss (Figure 9). Further, PD 144418 (Figure 9A), PB-28 (Figure 9D), and haloperidol (Figure 9F) restored spine density to levels that were not statistically different from those of neurons treated with mock-purified material.

**Figure 9:**
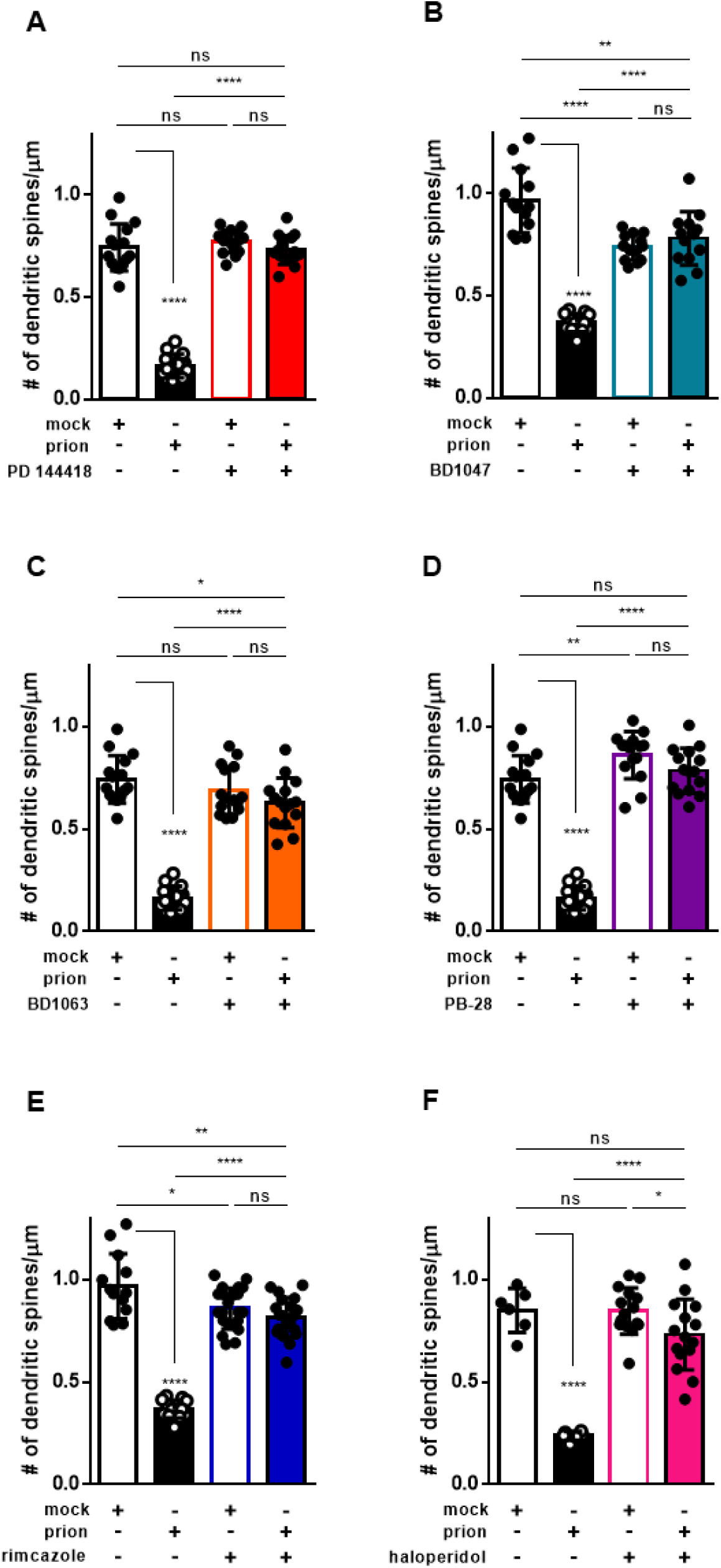
Sigma receptor ligands prevent prion induced synaptotoxicity. Following a 2 hour pre-incubation with the indicated compound, or DMSO vehicle, hippocampal neurons were exposed to PrP^Sc^ purified from the brains of RML-infected mice (prion), or to mock-purified material from uninfected brains (mock). Dendritic spine number was assayed 24 h later by imaging F-actin in spines using Alexa Fluor 488 conjugated phalloidin. Mean number of dendritic spines/μm ± SEM is plotted for the following treatments: **A)** 10 μM PD 144418; **B)** 0.5 μM BD1047; **C)** 1 μM BD1063; **D)** 0.1 μM PB-28; **E)** 0.5 μM Rimcazole; **F)** 2.5 μM haloperidol. All numerical values can be found in Supplementary Table 2. p < 0.0001 = ****; p < 0.001 = ***; p < 0.01 = **; p < 0.05 = *; ns = not significant using a two-tailed student’s t-test.

## Discussion

### Identification of sigma receptor ligands as anti-prion agents

Using the DBCA, we previously identified phenethyl piperidines as a novel class of anti-prion compound, capable of reducing the levels of PK resistant PrP in RML and 22L-infected ScN2a cells at low micromolar concentrations (20,21). To identify the target of these molecules relevant to their anti-prion effects we submitted JZ107, a structurally optimized phenethyl piperidine, and JZ103, an inactive analogue, to the PDSP. In this screen, the highest affinity receptors for JZ107 were σ_1_R and σ_2_R, both of which were highly expressed in N2a cells (Table 1). Based on this result, we tested known ligands for σ_1_R and σ_2_R, and identified ten novel anti-prion compounds: PD 144418, BD1047, BD1063, PB-28, rimcazole, haloperidol, SA 4503, ANAVEX2-73, (+)-pentazocine, and DTG (Figure 1; Table 2). Each of these molecules were found to significantly reduce PrP^Sc^ levels in ScN2a cells and to bind σ1R and σ2R with K_i_ values of 1.1 nM – 2.5 µM (Table 2).

### Exploration of the mechanism of action of sigma receptor ligands

σ1R and σ2R, originally thought to be a novel class of opioid receptors, were eventually shown to be a pharmacologically and molecularly distinct pair of receptors that reside primarily in ER membrane domains. Their endogenous ligands and physiological functions are uncertain (47–49). There is evidence that σ_1_R functions as a ligand-gated chaperone for other receptors, and that σ_2_R plays a role in cholesterol exchange between the ER and lysosomes; both receptors have an evolutionary relationship to sterol isomerases. Sigma receptors are being actively investigated as therapeutic targets for treatment of cancer, as well as neurological disorders, including neuropathic pain, amyotrophic lateral sclerosis, Alzheimer’s disease, and Parkinson’s disease (50).

Given their localization in the ER and their involvement in sterol metabolism, it seemed plausible that σ_1_R and σ_2_R could play a role in the formation or degradation of PrP^Sc^ in ScN2a cells, and thereby represent the molecular targets responsible for the anti-prion effects of the compounds identified in this study. However, a comparison of the affinity for σ_1_R or σ_2_R and the EC_50_ of these compounds against RML prions reveals that these parameters are not correlated (Figure 2). Further, dual knockout of the genes encoding σ_1_R and σ_2_R in ScN2a cells did not alter either basal levels of PK resistant PrP (Figure 3) or blunt the susceptibility of the cells to the anti-prion effects of the compounds (Figure 4). These results conclusively demonstrate that σ_1_R and σ_2_R are not the direct molecular targets responsible for the anti-prion effects of these compounds. This was surprising, given that we chose to test the anti-prion activity of these compounds based on the fact that they are well known ligands for the sigma receptors. However, σ_1_R and σ_2_R are known to be promiscuous in their ligand binding profiles, and binding to the sigma receptors does not rule out the possibility that these ligands may act via other molecular targets. To explore the possibility that these compounds may be acting though other, distinct, interaction partners, we applied a similar experimental approach to test five other target proteins identified by the PDSP (*Htr6*, *Drd4*, *Hrh1*, *Hrh3*, and *Chrm4*). However, similarly to what we observed with the sigma receptors, these additional target knockouts did not diminish the observed anti-prion effects (Figure 5).

### Other potential mechanisms of anti-prion activity

During the course of this study, the sigma receptors and their ligands were thrust into the spotlight for a potential role in the pathogenesis and treatment of SARS-CoV-2 infection (37). However, the affinity of these compounds for the sigma receptors did not correlate with their anti-viral activity (38). It was subsequently determined that the anti-viral effects were due to the ability of these compounds to induce phospholipidosis, a poorly understood phenomenon that often confounds drug discovery efforts. Because these findings included some of the compounds under examination in the present work, and because similar mechanisms could plausibly disrupt prion propagation, we next investigated phospholipidosis as a potential mechanism of action for these anti-prion compounds (Figure 6). While we found no correlation between phospholipidosis induction and the anti-prion effects of the molecules currently under investigation, we cannot rule out the possibility that phospholipidosis is a mechanism whereby some anti-prion compounds exert their effects, such as amiodarone, or that other kinds of disruptions of lysosomal or autophagosomal pathways contribute to reductions in PrP^Sc^.

We determined that none of the six most potent sigma receptor ligands identified here had a significant effect on the total levels or cellular localization of PrP^C^ (Figure 7; Supplementary Figure 1). This observation argues against an inhibitory mechanism dependent on reducing or redistributing the PrP^C^ substrate, as has been proposed for some anti-prion compounds (16,51). Although there were some variations in the potency of the compounds when tested on ScN2a cells infected with the RML and 22L strains, all of them significantly reduced PrP^Sc^ levels in cells infected with either strain, suggesting lack of a strong preference for particular PrP^Sc^ conformations (Figure 8).

A striking observation that may provide mechanistic insight is that all six of the most potent sigma receptor ligands, like JZ107 (20), significantly reduced PrP^Sc^-induced retraction of dendritic spines on cultured hippocampal neurons (Figure 9). We have previously demonstrated that spine retraction and decrements in synaptic transmission occur rapidly (< 12 hrs) after exposure to PrP^Sc^, and these events are entirely dependent on the expression of full length PrP^C^ by target neurons (22,45,52). We have also shown that the synaptotoxic effects of PrP^Sc^ depend on activation of an NMDA receptor/p38 MAPK-mediated signal transduction cascade, likely initiated by formation of PrP^Sc^ at the cell surface. The ability of the sigma receptor ligands to block this cascade could result from their effects on PrP^C^ or PrP^Sc^ at the neuronal plasma membrane, or on subsequent steps of the synaptotoxic signaling pathway.

### Therapeutic implications

The compounds identified here have several properties that make them attractive candidates as therapeutics for prion disease. In addition to their ability to inhibit prion propagation in ScN2a cells infected with multiple prion strains, the most potent compounds prevent prion-induced retraction of hippocampal neuron dendritic spines. This event is one of the earliest pathologies observed over the course of prion disease (53). Further, these compounds are all known to be blood brain barrier penetrant, and five of them, rimcazole, haloperidol, SA 4503, ANAVEX2-73, and (+)-pentazocine have a history of use in humans (49,54–62). The preclinical molecules SA 4503 and ANAVEX2-73, known respectively as cutamesine and blarcamesine, were well tolerated in phase II trials: SA 4503 for ischemic stroke (60), and ANAVEX2-73 for Alzheimer’s and Parkinson’s diseases (61). Despite the negative side effects associated with high and/or prolonged dosing with rimcazole, haloperidol, and (+)-pentazocine (57–59), all five molecules make excellent candidates for preclinical studies for the treatment of prion disease.

## Experimental Procedures

### Psychoactive Drug Screening Program

All information related to the PDSP, including detailed protocols, can be accessed at https://pdsp.unc.edu/pdspweb/. σ_1_R and σ_2_R binding assays were done using membrane fractions prepared from HEK 293T cells expressing the respective receptors. [^3^H]-Pentazocine was used to determine K_i_ values for σ_1_R and [^3^H]-DTG was used to determine K_i_ values for σ_2_R.

### Cell culture

For all experiments, N2a cells were grown in Opti-MEM (Gibco) supplemented with 10% FBS (Gemcell) and 100 U/ml each of penicillin/streptomycin (Gibco) in a humidified atmosphere of 5% CO_2_ at 37 °C. Chronic infection with RML or 22L prions was achieved by exposure of confluent cultures infected brain homogenate (1% final concentration) for 24 h before removal and subsequent passage.

### Drug treatment

Cells were incubated in the presence of each compound or DMSO vehicle (0.1% final DMSO concentration) for three days, before being split 1:5 and incubated for an additional four days in continued presence of compound or DMSO vehicle. Cells were then washed with PBS and lysed using cell lysis buffer containing 10 mM Tris pH 7.8, 100 mM NaCl, 0.5% NP-40, 0.5% Na deoxycholate and 0.1% SDS. All compounds were purchased from Tocris.

### MTT assay

Cells were washed 2x with PBS and incubated in the presence of 0.5 mg/ml MTT at 37 °C for 30 min. This solution was removed, and cells were incubated in DMSO at 37 ℃ for 10 min. Absorbance was then read at 570 nm using a Synergy H1 plate reader (BioTek). Signal was normalized to DMSO treatment and curves were fit by least squares regression using GraphPad software.

### Proteinase K treatment

100 μg of protein, as determined by BCA assay (Pierce), was exposed to 10 μg/ml proteinase K (Roche) in a final volume of 250 μL lysis buffer at 37 °C for 1 hour with shaking at 750 rpm. 30 µL of 10x protease inhibitor (Pierce) was added and samples were centrifuged at 21,130 x g for 1 hour at 4 °C. Supernatant was removed and pellets were resuspended in 1x Laemmli buffer (BioRad).

### Western blot

Protein samples were boiled in the presence of 1x Laemmli buffer (BioRad) and loaded into 12% Criterion TGX Precast Protein Gels and run at 200 V for 42 min. Proteins were transferred to PVDF membranes for 45 min at 115 V before gentle washing in 0.1% TBST and blocking in 5% non-fat milk in 0.1% TBST for 1 hour. The following primary antibodies were used: D18 (anti-PrP, made in house (63)); σ_1_R (B-5, Santa Cruz Biotechnology); σ_2_R (26444-1-AP, Proteintech); β-actin (AC-74, Millipore Sigma). All HRP-conjugated secondary antibodies were purchased from BioRad. Membranes were visualized using ECL (Millipore). Signal was normalized to DMSO treatment following quantification using ImageJ and plotted using GraphPad software.

### Immunofluorescence

For immunofluorescence staining of PrP^C^, cells were plated on coverslips during the passage following three days of treatment with compounds. Cells were washed with PBS and fixed using 4% paraformaldehyde in PBS pH 7.4 for 15 min at room temperature. Cells were then washed 3 x 5 min with PBS before incubation in 2% bovine serum albumen (BSA) and 20 mg/ml glycine in PBS with 0.1% Tween-20 for 30 min. Cells were then incubated in 10 ug/ml D18 for 1 hour at room temperature by inverting the coverslip over 30 μl of antibody solution spotted on parafilm, washed 3 x 5 min with PBS and incubated with Alexa Fluor-488 goat anti human secondary (Invitrogen) for 1 hour at room temperature. Cells were then washed 3 x 5 min with PBS and mounted on slides using mounting media with DAPI. Imaging was performed using a Zeiss Axio Observer Z1 microscope equipped with a digital camera (C10600/ORCA-R2 Hamamatsu Photonics). Images were taken using Zen software.

### CRISPR/Cas9 mediated gene disruption

We utilized a high-efficiency gene editing kit from Synthego (Gene Knockout Kit v2), which employs three chemically modified sgRNAs for each targeted locus (Rosa26 is targeted by a single guide): *Sigmar1* 5’-CCGUGUACAACUGUCUCUCC-3’, 5’-CCAGGAGAGACAGUUGUACA-3’, 5’- CAGGAGAGACAGUUGUACAC-3’; *Tmem97* 5’-UGCAGUUCAGCAACCUGUUG-3’,5’- ACUGUACCAACCUUUGAAGA-3’, 5’-CAUAUGCCUUCUUCAAAGGU-3’; *Hrh1* 5’- CUCACACAUCUUGUCUUCGG-3’, 5’-GGUUCUAAGUAGUAUCUCCC-3’, 5’- GAGCGCAAGCUACACACCGU-3’; *Hrh3* 5’- GCUCAUGGCGCUGCUCAUCG-3’, 5’- GCCUGCAGCCGCCGCCUCUC-3’, 5’- AUGGAGCGCGCGCCGCCCGA-3’; *Htr6* 5’- UUAGACGUGUUGCGCAGCGC-3’, 5’- GUGCGUGGUCAUCGUGCUGA-3’, 5’- AACAGUAGCACCCCAGCCUG-3’; *Chrm4* 5’- GACUGUGGUGGGUAACAUCC-3’, 5’- CUGCACACGCCAGGCUGAAC-3’, 5’-UGAUGAUGUAUAAGGUGUAA-3’; *Drd4* 5’- AGCAGCGCUACUGAGGACGG-3’, 5’-CUGGGGACUGGCGCCGGGCU-3’, 5’- GGGCGUGCUGCUCAUCGGCU-3’; Rosa26 5’- ACTCCAGTCTTTCTAGAAGA-3’. Recombinant Cas9 containing two nuclear localization signals (Synthego) is then mixed with sgRNAs to form ribonucleoprotein complexes that are then delivered by electroporation. This procedure allowed us to propagate gene-edited cells as a population, without the need for cloning. Electroporation was performed using the Amaxa Cell Line Nucleofector kit V (Lonza Bioscience) with the manufacturer’s settings for N2a cells. To evaluate knockout efficiency, genomic DNA was extracted and targeted regions were amplified using Q5 high fidelity polymerase (New England Biolabs) using the following primers: *Sigmar1*-F 5’- AAAGGCCAGAAGAGGGCATC-3’, *Sigmar1*-R 5’- CTGGGTGGATGTGAGTGCAT-3’, *Tmem97*-F 5’- TGCAGCTCTACAGTAGCATGT-3’, *Tmem97*-R 5’- GCTGGACACCAAACTGAGGT-3’, *Hrh1*-F 5’-ACTCCCAGTCTGACCACCAT-3’, *Hrh1*-R 5’- CGTGCTGGCCACATAATCCA-3’, *Hrh3*-F 5’-GACTCTGGCAGCGGACA-3’, *Hrh3*-R 5’- CAGATTCCGATACCAGCCCC-3’, *Htr6*-F 5’-AGCACATCACGTCGAAGGC-3’, *Htr6*-R 5’- AACCCTGTTTTGCCACCTAC-3’, *Chrm4*-F 5’-GTCTCCAGTTGGGTTCAGCA-3’, *Chrm4*-R 5’- GCGGGCTGGATAGGTGA-3’, *Drd4*-F 5’-AGCATTTCTCCCTCTGCCAA-3’, *Drd4*-R 5’- AGGCTCACCTCGGAGTAGAC-3’, Rosa26-F 5’-GAGGCGGATCACAAGCAATA-3’, Rosa26-R 5’- GGGAGGGGAGTGTTGCAATA-3’. Sanger sequencing was performed with the following primers: *Sigmar1* 5’-TCAAGCAGCTTGGCCAGTAGGGTAG-3’; *Tmem97* 5’- CAGTTGGAGCATGTGACTGCTGCC-3’; *Hrh1* 5’-TGCTGGCCACATAATCCATAGAG-3’; *Hrh3* 5’- TCTGGGGGCTTTACCCACGAGGAAGTCGGA-3’; *Htr6* 5’- GTCCAAAGCAGACAGAGGCCGCGAGCTAGC-3’; *Chrm4* 5’-GCGGGCTGGATAGGTGAGGG-3’; *Drd4* 5’-GCTCACCTCGGAGTAGACAAAG-3’; Rosa26 amplicons were sequenced with the forward primer listed above. Editing efficiency was determined with the Synthego Inference of CRISPR Edits (ICE) tool (https://ice.synthego.com) (36).

### RNA-seq and RT-PCR

Total RNA was extracted using the RNeasy mini plus extraction kit (Qiagen) following manufacturer’s instructions. RNA purity and concentration were measured using a nanodrop (Thermo). Library preparation and sequencing was performed by Novogene (Beijing, China). Genes with FPKM > 0.5 were considered expressed. For RT-qPCR analysis, 1 μg total RNA was converted to cDNA using the iScript cDNA synthesis kit (BioRad). RT-qPCR reactions were performed using Fast Sybr Master Mix (ABI) on a ViiA7 Real-Time PCR system. Relative mRNA expression was determined after normalization to the housekeeping gene *Actb*. The following primers were used: *Sigmar1*-F: 5’- GTCTGAGTACGTGCTGCTCTTC-3’, *Sigmar1*-R: 5’- GAAGACCTCACTTTTCGTGGTGC-3’, *Tmem97*-F: 5’- TCACGCTGTTCATCGACCTGCA-3’, *Tmem97*-R: 5’- GGAAGGACTTGAACCACACTGG-3’. *Prnp*-F: 5’- CAGCAACCAGAACAACTTCGTGC-3’, *Prnp*-R: 5’-CGCTCCATCATCTTCACATCGG-3’, *Actb*-F: 5’- CATTGCTGACAGGATGCAGAAGG-3’, *Actb*-R: 5’-TGCTGGAAGGTGGACAGTGAGG-3’. Relative fold changes were calculated using the delta-delta Ct method.

### LipidTox assay

Uninfected N2a cells were seeded in black 96-well glass bottom plates and incubated with 20 μM of compound or DMSO alone with 1x HCS LipidTOX™ Green neutral lipid stain (Invitrogen) for 24-48 hours. Cells were then rinsed with 3x with PBS before fixation with 4% paraformaldehyde in PBS with 10 μg/ml Hoechest 33342 (Tocris) for 30 min at RT. Cells were then washed 3x PBS before being read on a plate reader at the following wavelengths: ex/em 495/525 nm for LipidTox; 350/450 nm for Hoechest 33342. LipidTox signal was normalized to Hoechest 33342 signal for plotting. Fluorescence was also visualized with a Zeiss Axio Observer Z1 microscope using a 10x objective and Zen software.

### Dendritic spine retraction assay

Cultures of hippocampal neurons were performed as described previously (22). Neurons were preincubated for 2 h with compound (or DMSO vehicle for the controls), followed by treatment for 24 hr with 4.4 µg/ml of purified PrP^Sc^, or an equivalent volume of mock-purified control sample from non-infected brains. Neurons were then fixed in 4% paraformaldehyde and stained with Alexa Fluor 488- phalloidin (Thermo Fisher) to visualize actin in dendritic spines. Images were acquired using a Zeiss 700 confocal microscope with a 63x objective using Zen software. The number of spines was normalized to the measured length of the dendritic segment to give the number of spines/μm using ImageJ software. All animal studies were approved by the Boston University Chobanian & Avedisian School of Medicine Institutional Animal Care and Use Committee and performed in accordance to the United States Department of Agriculture Animal Welfare Act and the National Institutes of Health Policy on Humane Care and Use of Laboratory Animals.

### Prion purification

Prion purification was performed as described previously (46). Briefly, 200 μl of 10% (w/v) brain homogenate from a RML inoculated C57BL6/J mouse at the terminal phase of disease was treated with 100 μg/ml of pronase E (Sigma) for 30 min at 37 °C with shaking in a thermal mixer at 800 rpm. EDTA was added to a final concentration of 10 mM (pH 8) before adding sarkosyl in D-PBS and Benzonase to final concentrations of 2% (w/v) and 50 U/ml, respectively. After incubation at 37 °C for 10 min, samples were brought to 0.3% (w/v) NaPTA (pH 7.4) and incubated at 37 °C for 30 min. Samples were then mixed with iodixanol to a final concentration of 35% (w/v) adding enough additional NaPTA to maintain the final concentration of 0.3% (w/v). Samples were then centrifuged at 16, 100 x g for 90 min at room temperature. The upper layer was removed and filtered with a 0.45 μm pore size Durapore membrane Ultrafree-HV microcentrifuge filtration unit (Millipore). Samples were mixed with an equal volume of 0.3% NaPTA and 2% sarkosyl and incubated at 37 °C for 10 min before centrifugation for 90 min at 16, 100 x g. The supernatant was discarded and the pellet resuspended in D-PBS with 17.5% iodixanol and 0.1% sarkosyl. Following two washes, samples were resuspended in D-PBS with 0.1% (w/v) sarkosyl, pooled and stored as aliquots at −80 °C. Samples were analyzed by silver stain (per manufactures instruction; Pierce) and immunoblotting with and without PK digestion.

## Author contributions

Conducted experiments: RCCM, NTTL, MCQH; Designed experiments: RCCM, DAH; Contributed materials/methods: ABB; Project supervision: DAH; writing/editing: RCCM, DAH

## Acknowledgements

We would like to acknowledge the contributions of Dr. Thibaut Imberdis to early phases of this project. K_i_ determination was performed by the National Institute of Mental Health’s Psychoactive Drug Screening Program, Contract # HHSN-271-2018-00023-C (NIMH PDSP). The NIMH PDSP is directed by Dr. Bryan L. Roth at the University of North Carolina at Chapel Hill and Project Officer Jamie Driscoll at NIMH, Bethesda MD, USA. RML and 22L brain samples were generously provided by Drs. Byron Caughey and Brent Race at the National Institutes of Health. This project was funded by NIH- 5R01NS065244 to David A. Harris. Robert C.C. Mercer is supported by grants from the Department of Defense (W81XWH-21-1-0141) and the Creutzfeldt-Jakob Disease Foundation.

## Conflict of interest

The authors declare that they have no conflicts of interest with the contents of this article.

## Supplementary Figures

**Supplementary Figure 1: Western blots for Figure 7A**

Following treatment with the indicated compound, total PrP in cell lysates was analyzed by western blotting. **A)** PD 144418; **B)** BD1047; **C)** BD1063; **D)** PB-28; **E)** rimcazole; **F)** haloperidol.

**Supplementary Figure 2: Western blots for Figure 8A**

Following treatment with the indicated compound, PK-resistant PrP in cell lysates was analyzed by western blotting. **A)** PD 144418; **B)** BD1047; **C)** BD1063; **D)** PB-28; **E)** rimcazole; **F)** haloperidol.

**Supplementary Figure 3: Analysis of RML prion purification**

PrP_Sc_ purified from RML infected mouse brains, and mock-purified material from age-matched, uninfected brains was subjected to PK digestion and analysis by **A)** silver staining and **B)** western blot with the C-terminal anti prion antibody, D18.

## Notes

### Competing Interest Statement

The authors have declared no competing interest.

## References

1. Prusiner, S. B. (1998) Prions. Proc. Natl. Acad. Sci. USA 95, 13363–13383

2. Mercer, R. C., Daude, N., Dorosh, L., Fu, Z.-L., Mays, C. E., Gapeshina, H., Wohlgemuth, S. L., Acevedo-Morantes, C. Y., Yang, J., and Cashman, N. R. (2018) A novel Gerstmann-Sträussler-Scheinker disease mutation defines a precursor for amyloidogenic 8 kDa PrP fragments and reveals N-terminal structural changes shared by other GSS alleles. PLoS pathogens 14, e1006826

3. Medori, R., Tritschler, H.-J., LeBlanc, A., Villare, F., Manetto, V., Chen, H. Y., Xue, R., Leal, S., Montagna, P., and Cortelli, P. (1992) Fatal familial insomnia, a prion disease with a mutation at codon 178 of the prion protein gene. New England Journal of Medicine 326, 444–449

4. Mercer, R. C., McDonald, A. J., Bove-Fenderson, E., Fang, C., Wu, B., and Harris, D. A. (2018) Prion diseases. in The Molecular and Cellular Basis of Neurodegenerative Diseases, Elsevier. pp 23–56

5. Kraus, A., Hoyt, F., Schwartz, C. L., Hansen, B., Artikis, E., Hughson, A. G., Raymond, G. J., Race, B., Baron, G. S., and Caughey, B. (2021) High-resolution structure and strain comparison of infectious mammalian prions. Molecular Cell 81, 4540–4551. e4546

6. Manka, S. W., Zhang, W., Wenborn, A., Betts, J., Joiner, S., Saibil, H. R., Collinge, J., and Wadsworth, J. D. (2022) 2.7 Å cryo-EM structure of ex vivo RML prion fibrils. Nature communications 13, 1–11

7. Manka, S. W., Wenborn, A., Betts, J., Joiner, S., Saibil, H. R., Collinge, J., and Wadsworth, J. D. (2023) A structural basis for prion strain diversity. Nature Chemical Biology, 1-7

8. Hoyt, F., Alam, P., Artikis, E., Schwartz, C. L., Hughson, A. G., Race, B., Baune, C., Raymond, G. J., Baron, G. S., and Kraus, A. (2022) Cryo-EM of prion strains from the same genotype of host identifies conformational determinants. PLoS pathogens 18, e1010947

9. Collinge, J., Gorham, M., Hudson, F., Kennedy, A., Keogh, G., Pal, S., Rossor, M., Rudge, P., Siddique, D., and Spyer, M. (2009) Safety and efficacy of quinacrine in human prion disease (PRION-1 study): a patient-preference trial. The Lancet Neurology 8, 334–344

10. Haïk, S., Marcon, G., Mallet, A., Tettamanti, M., Welaratne, A., Giaccone, G., Azimi, S., Pietrini, V., Fabreguettes, J.-R., and Imperiale, D. (2014) Doxycycline in Creutzfeldt-Jakob disease: a phase 2, randomised, double-blind, placebo-controlled trial. The Lancet Neurology 13, 150–158

11. Berry, D. B., Lu, D., Geva, M., Watts, J. C., Bhardwaj, S., Oehler, A., Renslo, A. R., DeArmond, S. J., Prusiner, S. B., and Giles, K. (2013) Drug resistance confounding prion therapeutics. Proceedings of the National Academy of Sciences 110, E4160–E4169

12. Cronier, S., Beringue, V., Bellon, A., Peyrin, J.-M., and Laude, H. (2007) Prion strain-and species-dependent effects of antiprion molecules in primary neuronal cultures. Journal of virology 81, 13794–13800

13. Kocisko, D. A., Engel, A. L., Harbuck, K., Arnold, K. M., Olsen, E. A., Raymond, L. D., Vilette, D., and Caughey, B. (2005) Comparison of protease-resistant prion protein inhibitors in cell cultures infected with two strains of mouse and sheep scrapie. Neuroscience letters 388, 106–111

14. Baral, P. K., Swayampakula, M., Rout, M. K., Kav, N. N., Spyracopoulos, L., Aguzzi, A., and James, M. N. (2014) Structural basis of prion inhibition by phenothiazine compounds. Structure 22, 291–303

15. Nicoll, A. J., Trevitt, C. R., Tattum, M. H., Risse, E., Quarterman, E., Ibarra, A. A., Wright, C., Jackson, G. S., Sessions, R. B., and Farrow, M. (2010) Pharmacological chaperone for the structured domain of human prion protein. Proceedings of the National Academy of Sciences 107, 17610–17615

16. Masone, A., Zucchelli, C., Caruso, E., Lavigna, G., Eraña, H., Giachin, G., Tapella, L., Comerio, L., Restelli, E., and Raimondi, I. (2023) A tetracationic porphyrin with dual anti-prion activity. iScience

17. Sigurdson, C. J., Nilsson, K. P. R., Hornemann, S., Manco, G., Polymenidou, M., Schwarz, P., Leclerc, M., Hammarström, P., Wüthrich, K., and Aguzzi, A. (2007) Prion strain discrimination using luminescent conjugated polymers. Nature methods 4, 1023–1030

18. Margalith, I., Suter, C., Ballmer, B., Schwarz, P., Tiberi, C., Sonati, T., Falsig, J., Nyström, S., Hammarström, P., and Åslund, A. (2012) Polythiophenes inhibit prion propagation by stabilizing prion protein (PrP) aggregates. Journal of Biological Chemistry 287, 18872–18887

19. Poncet-Montange, G., St Martin, S. J., Bogatova, O. V., Prusiner, S. B., Shoichet, B. K., and Ghaemmaghami, S. (2011) A survey of antiprion compounds reveals the prevalence of non-PrP molecular targets. J. Biol. Chem. 286, 27718–27728

20. Imberdis, T., Heeres, J. T., Yueh, H., Fang, C., Zhen, J., Rich, C. B., Glicksman, M., Beeler, A., and Harris, D. A. (2016) Identification of anti-prion compounds using a novel cellular assay. J. Biol. Chem. 291, 26164–26176

21. Massignan, T., Stewart, R. S., Biasini, E., Solomon, I. H., Bonetto, V., Chiesa, R., and Harris, D. A. (2010) A novel, drug-based, cellular assay for the activity of neurotoxic mutants of the prion protein. Journal of Biological Chemistry 285, 7752–7765

22. Fang, C., Imberdis, T., Garza, M. C., Wille, H., and Harris, D. A. (2016) A Neuronal Culture System to Detect Prion Synaptotoxicity. PLoS pathogens 12, e1005623

23. Mercer, R. C., and Harris, D. A. (2019) Identification of anti-prion drugs and targets using toxicity-based assays. Current opinion in pharmacology 44, 20–27

24. Besnard, J., Ruda, G. F., Setola, V., Abecassis, K., Rodriguiz, R. M., Huang, X.-P., Norval, S., Sassano, M. F., Shin, A. I., and Webster, L. A. (2012) Automated design of ligands to polypharmacological profiles. Nature 492, 215–220

25. Akunne, H., Whetzel, S., Wiley, J., Corbin, A., Ninteman, F., Tecle, H., Pei, Y., Pugsley, T., and Heffner, T. (1997) The pharmacology of the novel and selective sigma ligand, PD 144418. Neuropharmacology 36, 51–62

26. Matsumoto, R. R., Bowen, W. D., Tom, M. A., Truong, D. D., and De Costa, B. R. (1995) Characterization of two novel σ receptor ligands: antidystonic effects in rats suggest σ receptor antagonism. Eur. J. Pharmacol. 280, 301–310

27. Colabufo, N. A., Abate, C., Contino, M., Inglese, C., Ferorelli, S., Berardi, F., and Perrone, R. (2008) Tritium radiolabelling of PB28, a potent sigma-2 receptor ligand: pharmacokinetic and pharmacodynamic characterization. Bioorg. Med. Chem. Lett. 18, 2183–2187

28. Ferris, R., Tang, F., Chang, K.-J., and Russell, A. (1986) Evidence that the potential antipsychotic agent rimcazole (BW 234U) is a specific, competitive antagonist of sigma sites in brain. Life sciences 38, 2329–2337

29. Tam, S. W., and Cook, L. (1984) Sigma opiates and certain antipsychotic drugs mutually inhibit (+)-[3H] SKF 10,047 and [3H] haloperidol binding in guinea pig brain membranes. Proceedings of the National Academy of Sciences 81, 5618–5621

30. Matsuno, K., Nakazawa, M., Okamoto, K., Kawashima, Y., and Mita, S. (1996) Binding properties of SA4503, a novel and selective σ1 receptor agonist. European journal of pharmacology 306, 271–279

31. Villard, V., Espallergues, J., Keller, E., Vamvakides, A., and Maurice, T. (2011) Anti-amnesic and neuroprotective potentials of the mixed muscarinic receptor/sigma1 (σ1) ligand ANAVEX2-73, a novel aminotetrahydrofuran derivative. Journal of Psychopharmacology 25, 1101–1117

32. Su, T.-P. (1982) Evidence for sigma opioid receptor: binding of [3H] SKF-10047 to etorphine- inaccessible sites in guinea-pig brain. Journal of Pharmacology and Experimental Therapeutics 223, 284–290

33. Weber, E., Sonders, M., Quarum, M., McLean, S., Pou, S., and Keana, J. (1986) 1, 3-Di (2-[5-3H] tolyl) guanidine: a selective ligand that labels sigma-type receptors for psychotomimetic opiates and antipsychotic drugs. Proceedings of the National Academy of Sciences 83, 8784–8788

34. Gewirtz, G. R., Gorman, J. M., Volavka, J., Macaluso, J., Gribkoff, G., Taylor, D. P., and Borison, R. (1994) BMY 14802, a sigma receptor ligand for the treatment of schizophrenia. Neuropsychopharmacology 10, 37–40

35. Mahal, S. P., Baker, C. A., Demczyk, C. A., Smith, E. W., Julius, C., and Weissmann, C. (2007) Prion strain discrimination in cell culture: the cell panel assay. Proceedings of the National Academy of Sciences 104, 20908–20913

36. Conant, D., Hsiau, T., Rossi, N., Oki, J., Maures, T., Waite, K., Yang, J., Joshi, S., Kelso, R., and Holden, K. (2022) Inference of CRISPR edits from Sanger trace data. The CRISPR Journal 5, 123–130

37. Gordon, D. E., Jang, G. M., Bouhaddou, M., Xu, J., Obernier, K., White, K. M., O’Meara, M. J., Rezelj, V. V., Guo, J. Z., and Swaney, D. L. (2020) A SARS-CoV-2 protein interaction map reveals targets for drug repurposing. Nature 583, 459–468

38. Tummino, T. A., Rezelj, V. V., Fischer, B., Fischer, A., O’meara, M. J., Monel, B., Vallet, T., White, K. M., Zhang, Z., and Alon, A. (2021) Drug-induced phospholipidosis confounds drug repurposing for SARS-CoV-2. Science 373, 541–547

39. Anderson, N., and Borlak, J. (2006) Drug-induced phospholipidosis. FEBS letters 580, 5533–5540

40. Shoup, D., and Priola, S. A. (2023) Cell biology of prion strains in vivo and in vitro. Cell and Tissue Research 392, 269–283

41. Klingenstein, R., Löber, S., Kujala, P., Godsave, S., Leliveld, S. R., Gmeiner, P., Peters, P. J., and Korth, C. (2006) Tricyclic antidepressants, quinacrine and a novel, synthetic chimera thereof clear prions by destabilizing detergent-resistant membrane compartments. Journal of neurochemistry 98, 748–759

42. Monassier, L., and Bousquet, P. (2002) Sigma receptors: from discovery to highlights of their implications in the cardiovascular system. Fundamental & clinical pharmacology 16, 1–8

43. Nioi, P., Perry, B. K., Wang, E.-J., Gu, Y.-Z., and Snyder, R. D. (2007) In vitro detection of drug-induced phospholipidosis using gene expression and fluorescent phospholipid–based methodologies. Toxicological sciences 99, 162–173

44. Kawasaki, Y., Kawagoe, K., Chen, C.-j., Teruya, K., Sakasegawa, Y., and Doh-Ura, K. (2007) Orally administered amyloidophilic compound is effective in prolonging the incubation periods of animals cerebrally infected with prion diseases in a prion strain-dependent manner. Journal of virology 81, 12889–12898

45. Fang, C., Wu, B., Le, N. T. T., Imberdis, T., Mercer, R. C. C., and Harris, D. A. (2018) Prions activate a p38 MAPK synaptotoxic signaling pathway. PLoS pathogens 14, e1007283

46. Wenborn, A., Terry, C., Gros, N., Joiner, S., D’Castro, L., Panico, S., Sells, J., Cronier, S., Linehan, J. M., and Brandner, S. (2015) A novel and rapid method for obtaining high titre intact prion strains from mammalian brain. Scientific Reports 5, 1–13

47. Schmidt, H. R., Zheng, S., Gurpinar, E., Koehl, A., Manglik, A., and Kruse, A. C. (2016) Crystal structure of the human σ1 receptor. Nature 532, 527–530

48. Alon, A., Schmidt, H. R., Wood, M. D., Sahn, J. J., Martin, S. F., and Kruse, A. C. (2017) Identification of the gene that codes for the σ2 receptor. Proceedings of the National Academy of Sciences 114, 7160–7165

49. Alon, A., Lyu, J., Braz, J. M., Tummino, T. A., Craik, V., O’Meara, M. J., Webb, C. M., Radchenko, D. S., Moroz, Y. S., and Huang, X.-P. (2021) Structures of the σ2 receptor enable docking for bioactive ligand discovery. Nature 600, 759–764

50. Schmidt, H. R., and Kruse, A. C. (2019) The molecular function of σ receptors: past, present, and future. Trends in pharmacological sciences 40, 636–654

51. Shyng, S.-L., Lehmann, S., Moulder, K. L., and Harris, D. A. (1995) Sulfated glycans stimulate endocytosis of the cellular isoform of the prion protein, PrPC, in cultured cells. Journal of Biological Chemistry 270, 30221–30229

52. Mercer, R. C., and Harris, D. A. (2022) Mechanisms of prion-induced toxicity. Cell and Tissue Research, 1–16

53. Fuhrmann, M., Mitteregger, G., Kretzschmar, H., and Herms, J. (2007) Dendritic pathology in prion disease starts at the synaptic spine. The Journal of neuroscience : the official journal of the Society for Neuroscience 27, 6224–6233

54. Lever, J. R., Miller, D. K., Fergason-Cantrell, E. A., Green, C. L., Watkinson, L. D., Carmack, T. L., and Lever, S. Z. (2014) Relationship between cerebral sigma-1 receptor occupancy and attenuation of cocaine’s motor stimulatory effects in mice by PD144418. Journal of Pharmacology and Experimental Therapeutics 351, 153–163

55. Urani, A., Roman, F. J., Phan, V.-L., Su, T.-P., and Maurice, T. (2001) The antidepressant-like effect induced by ς1-receptor agonists and neuroactive steroids in mice submitted to the forced swimming test. Journal of Pharmacology and Experimental Therapeutics 298, 1269–1279

56. Sabino, V., Cottone, P., Zhao, Y., Iyer, M. R., Steardo, L., Rice, K. C., Conti, B., Koob, G. F., and Zorrilla, E. P. (2009) The σ-receptor antagonist BD-1063 decreases ethanol intake and reinforcement in animal models of excessive drinking. Neuropsychopharmacology 34, 1482–1493

57. Gilmore, D. L., Liu, Y., and Matsumoto, R. R. (2004) Review of the pharmacological and clinical profile of rimcazole. CNS drug reviews 10, 1–22

58. Kornhuber, J., Schultz, A., Wiltfang, J., Meineke, I., Gleiter, C. H., Zöchling, R., Boissl, K.-W., Leblhuber, F., and Riederer, P. (1999) Persistence of haloperidol in human brain tissue. American Journal of Psychiatry 156, 885–890

59. Beach, S. R., Gross, A. F., Hartney, K. E., Taylor, J. B., and Rundell, J. R. (2020) Intravenous haloperidol: A systematic review of side effects and recommendations for clinical use. General Hospital Psychiatry 67, 42–50

60. Urfer, R., Moebius, H. J., Skoloudik, D., Santamarina, E., Sato, W., Mita, S., and Muir, K. W. (2014) Phase II trial of the Sigma-1 receptor agonist cutamesine (SA4503) for recovery enhancement after acute ischemic stroke. Stroke 45, 3304–3310

61. Hampel, H., Williams, C., Etcheto, A., Goodsaid, F., Parmentier, F., Sallantin, J., Kaufmann, W. E., Missling, C. U., and Afshar, M. (2020) A precision medicine framework using artificial intelligence for the identification and confirmation of genomic biomarkers of response to an Alzheimer’s disease therapy: analysis of the blarcamesine (ANAVEX2-73) Phase 2a clinical study. Alzheimer’s & Dementia: Translational Research & Clinical Interventions 6, e12013

62. Suzuki, T., Oshimi, M., Tomono, K., Hanano, M., and Watanabe, J. (2002) Investigation of transport mechanism of pentazocine across the blood-brain barrier using the in situ rat brain perfusion technique. Journal of pharmaceutical sciences 91, 2346–2353

63. Safar, J. G., Scott, M., Monaghan, J., Deering, C., Didorenko, S., Vergara, J., Ball, H., Legname, G., Leclerc, E., and Solforosi, L. (2002) Measuring prions causing bovine spongiform encephalopathy or chronic wasting disease by immunoassays and transgenic mice. Nature biotechnology 20, 1147–1150

64. Kronenberg, E., Weber, F., Brune, S., Schepmann, D., Almansa, C., Friedland, K., Laurini, E., Pricl, S., and Wünsch, B. (2019) Synthesis and Structure–Affinity Relationships of Spirocyclic Benzopyrans with Exocyclic Amino Moiety. Journal of Medicinal Chemistry 62, 4204–4217

65. Azzariti, A., Colabufo, N. A., Berardi, F., Porcelli, L., Niso, M., Simone, G. M., Perrone, R., and Paradiso, A. (2006) Cyclohexylpiperazine derivative PB28, a σ2 agonist and σ1 antagonist receptor, inhibits cell growth, modulates P-glycoprotein, and synergizes with anthracyclines in breast cancer. Molecular cancer therapeutics 5, 1807–1816

66. Yano, H., Bonifazi, A., Xu, M., Guthrie, D. A., Schneck, S. N., Abramyan, A. M., Fant, A. D., Hong, W. C., Newman, A. H., and Shi, L. (2018) Pharmacological profiling of sigma 1 receptor ligands by novel receptor homomer assays. Neuropharmacology 133, 264–275

67. Hong, W. C. (2020) Distinct regulation of σ1 receptor multimerization by its agonists and antagonists in transfected cells and rat liver membranes. Journal of Pharmacology and Experimental Therapeutics 373, 290–301

68. Paquette, M. A., Foley, K., Brudney, E. G., Meshul, C. K., Johnson, S. W., and Berger, S. P. (2009) The sigma-1 antagonist BMY-14802 inhibits L-DOPA-induced abnormal involuntary movements by a WAY-100635-sensitive mechanism. Psychopharmacology 204, 743–754

69. Maurice, T., and Su, T.-P. (2009) The pharmacology of sigma-1 receptors. Pharmacology & therapeutics 124, 195–206

70. Martin, P. M., Ola, M. S., Agarwal, N., Ganapathy, V., and Smith, S. B. (2004) The sigma receptor ligand (+)− pentazocine prevents apoptotic retinal ganglion cell death induced in vitro by homocysteine and glutamate. Molecular brain research 123, 66–75

