## Supplementary figures and images for "Sigma receptor ligands are potent anti-prion compounds that act independently of sigma receptor binding"

### Supplemental Figure 1

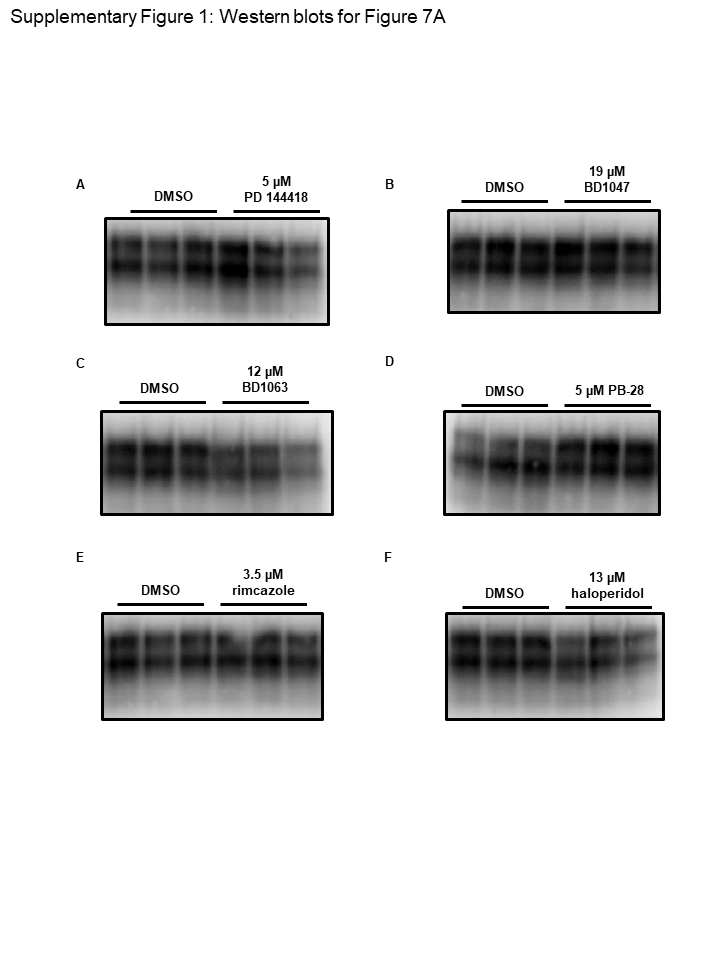

### Supplemental Figure 2

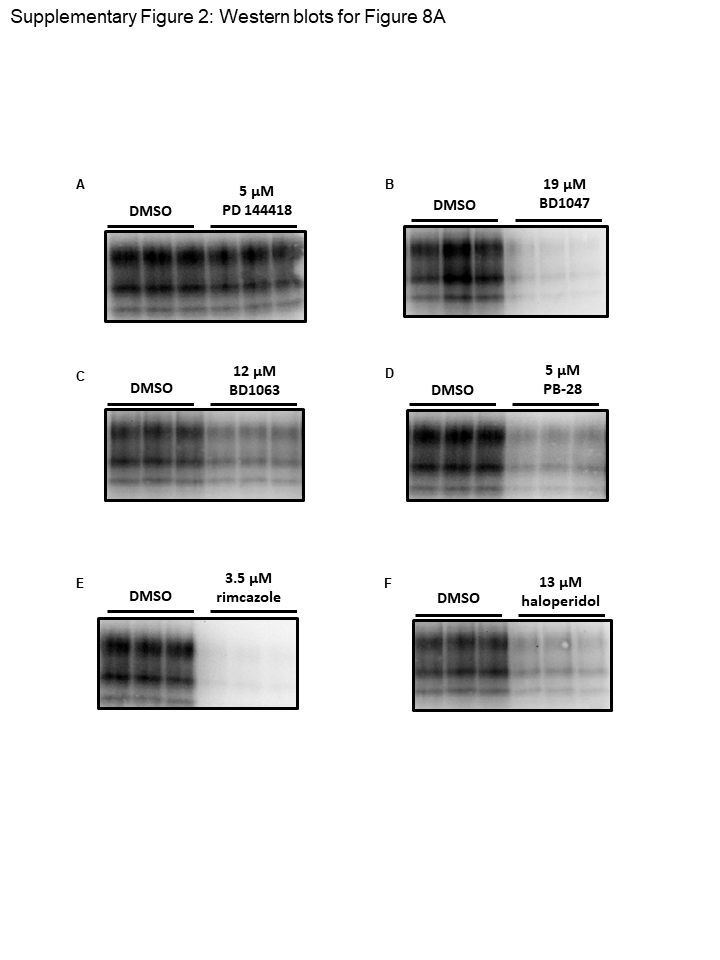

### Supplemental Figure 3

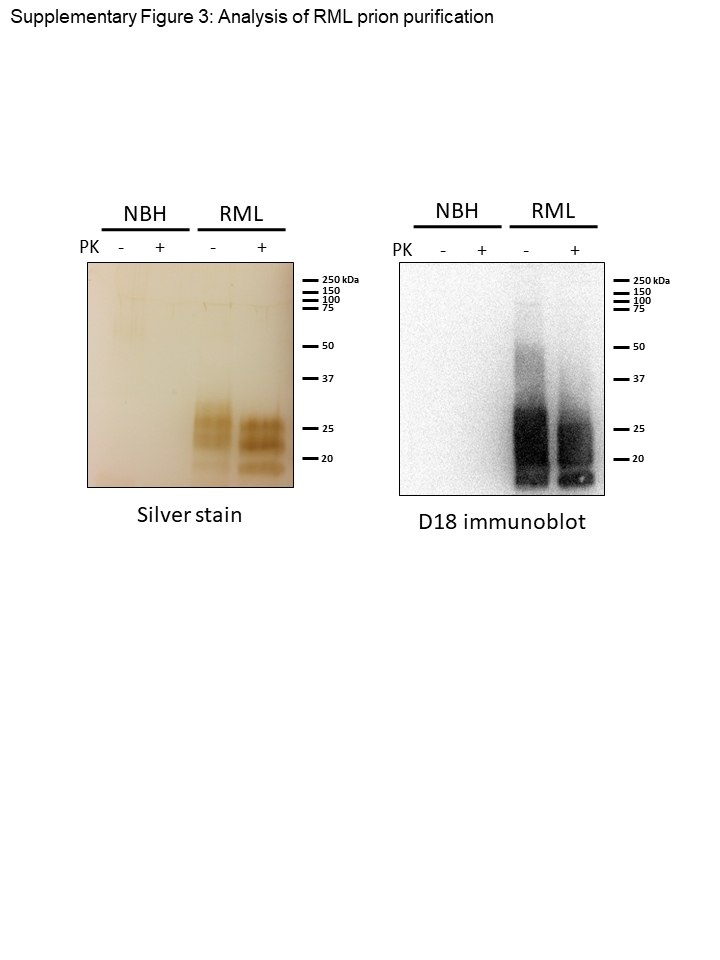
